# Integrative analysis of taste genetics and the dental plaque microbiome in early childhood caries

**DOI:** 10.1101/2025.03.10.642456

**Authors:** Mohd Wasif Khan, Vivianne Cruz de Jesus, Betty-Anne Mittermuller, Robert J. Schroth, Pingzhao Hu, Prashen Chelikani

## Abstract

Early childhood caries (ECC) is a multifactorial disease mainly caused by the oral microbiome; however, it is also influenced by host genetics and environmental factors. The combinatorial analysis of these multiple factors influencing ECC susceptibility requires further research. This study investigated the interplay between genetic variants in taste-related genes and the microbiome in ECC, targeting taste genes because of their role in taste preference and potential interactions with oral fungi and bacteria. Using a case-control design involving 538 children, we obtained dental plaque microbiome profiles and genetic variants across 55 candidate genes through next-generation sequencing. Our association analysis for taste genetics and ECC outcome used the socioeconomic factor index (SEFI) and rural-urban status as confounders. We observed a few taste gene variants associated with ECC and microbial diversity. However, no specific association was observed between the variants and cariogenic species. Furthermore, our analysis indicated that Streptococcus mutans is a partial mediator between these genetic variants and ECC outcomes. Machine learning models integrating microbiome, genetics, and covariates achieved robust ECC vs. caries-free classification (AUROC = 0.96), with Streptococcus mutans, rural-urban status, a bitter taste receptor variant, Candida dubliniensis, and SEFI as the top ECC-associated factors. Our findings highlight the association between host genetics and the oral microbiome, underscoring the need for multiomics approaches in ECC risk assessment.

**Highlights:**

- *S. mutans*, *C. dubliniensis*, and taste genetic variants are associated with ECC.
- Rural-urban status and SEFI score are among the top social markers for ECC prediction.
- Taste-related genetic factors modulate the composition of dental plaque microbiome.

## Introduction

Early childhood caries (ECC) refers to dental caries that affect children under 72 months of age (AAPD, 2024). It is a condition of the primary dentition. The high prevalence of ECC poses a significant socioeconomic burden on both families and society (Pierce et al., 2019; Tinanoff et al., 2019). Although microorganisms in the oral cavity, particularly in dental plaque, play a major role in ECC, numerous studies have highlighted additional factors, including dietary habits, oral hygiene practices, and host-related factors such as salivary composition, immune response, and tooth susceptibility (Baker et al., 2021; Opal et al., 2015). The etiology of ECC is complex; however, many of these contributing factors are closely linked to host genetics.

ECC is associated with dysbiosis of the oral microbiome, and diverse microbial species influence the progression of the disease. While *Streptococcus mutans* is considered a key cariogenic species, other species of genera, such as *Veillonella, Prevotella, Scardovia,* and *Candida,* consistently appear to be highly abundant in the oral microbiome (V. H. K. Lee et al., 2025; Tanner et al., 2011; Wang et al., 2017). Furthermore, the role of socioeconomic and behavioral factors in shaping the oral microbiome has gained a lot of attention recently (M. W. Khan, de Jesus, et al., 2024; V. H. K. Lee et al., 2025).

ECC is a complex health issue with a strong role of socioeconomic, behavioral and environmental factors. Numerous studies have identified a strong association between low socioeconomic status (SES) and ethnicity-deprived neighborhoods (Seow, 2012; Willems et al., 2005). Furthermore, behavioral factors such as poor feeding habits, infrequent tooth brushing, and lack of fluoride exposure significantly contribute to ECC development (M. W. Khan, de Jesus, et al., 2024; S. Y. Khan et al., 2023).

Host genetics plays a significant role in caries, with nearly half of the prevalence attributed to genetic factors (Haworth et al., 2020). Genetic predisposition has been linked to increased susceptibility to ECC, with specific genetic markers associated with this condition (Morrison et al., 2016; Olatosi et al., 2021; Shungin et al., 2019; Stanley et al., 2014). Twin studies examining host genetic influence have revealed higher similarities in the oral microbiome of monozygotic twins than in dizygotic twins, with more than 50% heritability of microbiome phenotypes (Demmitt et al., 2017). Additional studies have demonstrated a connection between host genetics and the oral microbiome (Blekhman et al., 2015; de Jesus et al., 2022; Gomez et al., 2017; Hashim et al., 2021; Kolde et al., 2018; Liu et al., 2021). Recent metagenome-genome association studies have identified several genomic loci associated with the oral microbiome (Liu et al., 2021), suggesting an indirect role of host genetics in the etiology of ECC mediated by microbiome interactions.

Recent studies have suggested that taste perception, mediated by T1R2 and T1R3 receptors for sweet and umami tastes and T2R receptors for bitter taste, may influence the oral microbiome and potentially affect caries susceptibility (Cattaneo et al., 2019; Chandrashekar et al., 2000; Chen et al., 2011; Chisini et al., 2021; de Jesus et al., 2022; Holla et al., 2015; Shrestha et al., 2024). Notably, single nucleotide variations (SNVs) in *TAS1R2* and *TAS2R38* have been linked to an increased risk of caries and the prevalence of specific bacterial genera (Cattaneo et al., 2019; de Jesus, Singh, et al., 2021; Kulkarni et al., 2013).

Despite these findings, the influence of host taste genetics on ECC and the associated microbiome remains largely underexplored. To address this, we aimed to utilize next-generation sequencing (NGS) technology, which can capture novel genetic variants across diverse populations. Additionally, we sought to identify the empirical changes in microbial profiles driven by these genetic variations. Research exploring the relationship between taste genetics and the oral microbiome is limited.

This study explored the relationship between the dental plaque microbiome, taste genetics, and ECC, addressing the following key questions. First, we investigated whether variants in taste and taste-related genes (referred to as “taste genes”) are associated with ECC and caries-free (CF) outcomes. Second, we examined whether these genetic variants influence the microbial composition of dental plaque, thereby mediating ECC outcomes through host genetics. Further, we compared the relative contributions of microbial and genetic factors using machine learning models. Third, we analyzed the role of the socioeconomic factor index (SEFI) and rural-urban status as confounders. The study design is shown in **Figure 1**. Our results suggest that certain taste gene variants are associated with ECC, and a variant in the bitter taste receptor *TAS2R60* influences the abundance of multiple dental plaque microbial species. Understanding these associations can contribute to deeper insights into disease etiology, children’s susceptibility to ECC, and strategies for risk assessment.

**Figure 1:**
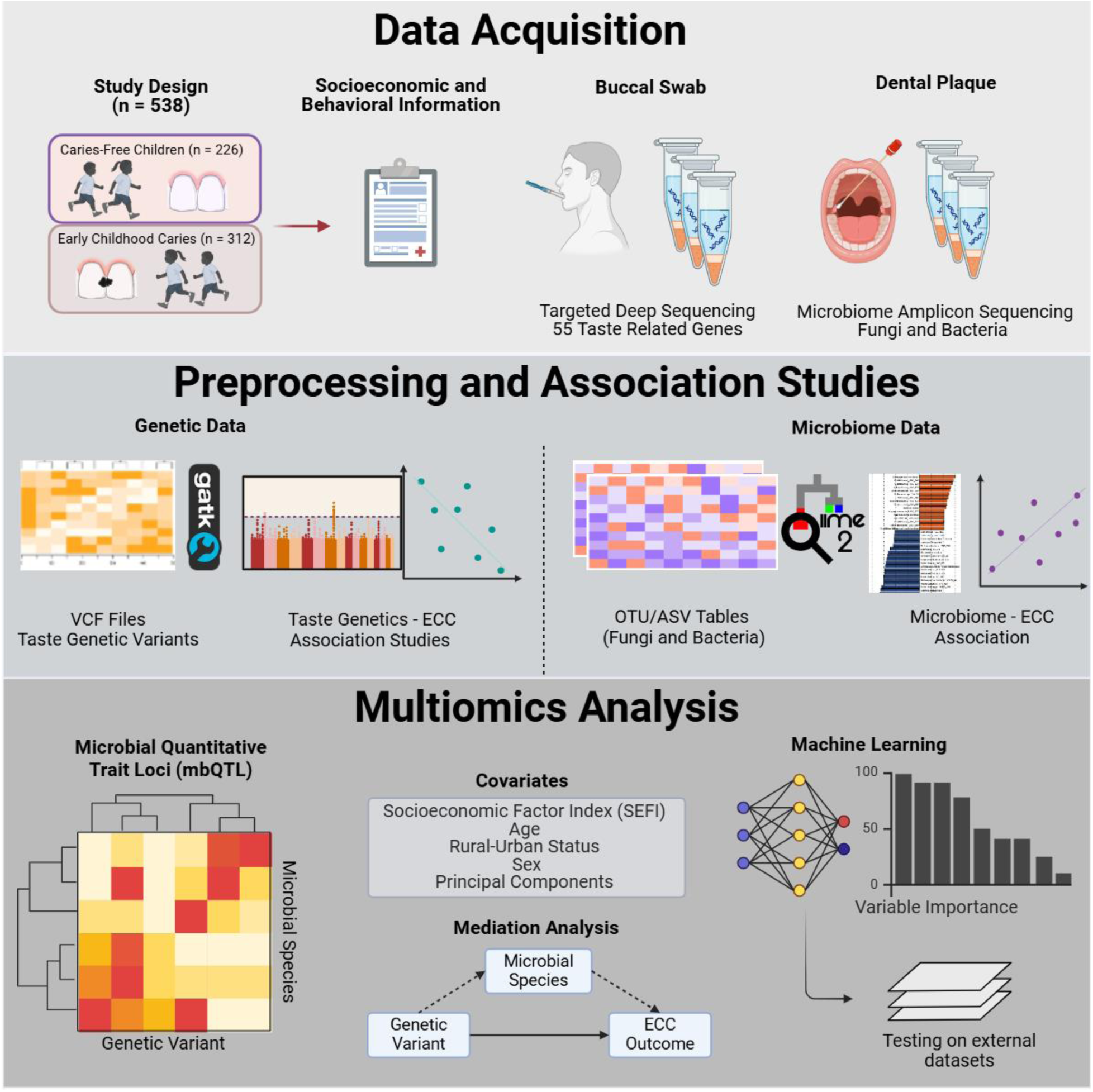
Overview of the study design and workflow for the combined microbial and taste genetic data analysis.

## Results

### Social demographical variables included in the study

Initially, 16 variables were selected for this study, as outlined in our previous publication (M. W. Khan, de Jesus, et al., 2024). However, based on their relevance and association with ECC, only age, sex, place of residence (rural-urban status), SEFI score, and ECC status were retained. The distributions of the selected variables are presented in **Table 1**. Age, rural-urban status, and SEFI score showed significant associations with ECC. These variables, along with the first five principal components (PCs) derived from the quality-filtered genetic data obtained in this study, were included as covariates in all subsequent statistical models. The first five PCs accounted for 73% of the total genetic variation in the dataset (**Figure S1**). A conceptual framework representing the relationships among covariates, independent variables, and outcomes of the study is presented in **Figure S2A**. Notably, the first PC effectively captured batch information with a coefficient of 0.81 (**Figure S2B**). Therefore, PC1 was used as a proxy for batch effects, and the batch variable was not included as a separate covariate.

**Table 1:** Descriptive statistics for the host variables used in the study.

| Variable | N | CF, n = 226<sup>1</sup> | ECC, n = 312<sup>1</sup> | p value<sup>2</sup> |
| --- | --- | --- | --- | --- |
| Sex | 538 |  |  | >0.9 |
| Female (0) |  | 114 (50%) | 157 (50%) |  |
| Male (1) |  | 112 (50%) | 155 (50%) |  |
| **Age** (months) | 538 | 43 (28, 55) | 49 (36, 58) | <0.001 |
| Urban_status | 534 |  |  | <0.001 |
| Rural (0) |  | 8 (3.6%) | 156 (50%) |  |
| Urban (1) |  | 216 (96%) | 154 (50%) |  |
| SEFI_score<br>(Socioeconomic Factor<br>Index score) | 498 | -0.06 (-0.67, 0.71) | 0.87 (0.04, 1.89) | <0.001 |

n, Number of non-missing entries for each variable
CF, Caries-free
ECC, Early childhood caries
<sup>1</sup>n (%); Median (IQR)
<sup>2</sup>Pearson's Chi-squared test; Wilcoxon rank sum test; Fisher's exact test

### Gene and genetic variant association testing with ECC

The sequencing data achieved a minimum depth of 30× for 95.4% of the reads across the 55 selected genes, covering an average of 99.4% of their exonic regions. Mapping these reads to the reference sequence for the selected genes using the standard GATK pipeline (McKenna et al., 2010), combined with initial quality control using VCFtools, identified 6,843 variants across 538 individuals. Subsequent quality control using the PLINK tool retained 696 genetic variants (Purcell et al., 2007), including single nucleotide variants (SNVs) and insertions/deletions (InDels), which were further tested for their association with ECC status.

The association of variants with ECC status was tested for different combinations of covariates (**Figure S3, S4, and S5**). These results show that while rural-urban status and SEFI score improve the model performance by decreasing the genomic inflation and Akaike Information Criterion (AIC) and Bayesian Information Criterion (BIC) values, the inclusion of the batch effect does not. Based on this optimization, the final model included the following covariates: age, sex, urban-rural status, SEFI score, and first five PCs. Using a *q*-value (Benjamini-Hochberg adjusted *p*-values) threshold of 0.05, three variants were significantly associated with ECC in the dominant model (**Figure 2A** and **2B**, **Table 2**): rs111819661 in *SCNN1D*, rs35195910 in *TAS2R60*, and rs2305645 in *PLCB2*. These variants also displayed significant deviations from the expected *p*-values in the quantile-quantile (QQ) plot (**Figure 2A**). In the additive model, only one variant, rs111819661 in *SCNN1D*, remained significant, and no association was identified in the recessive model (**Table 2** and **Figure S6**). The genomic inflation factors for both the dominant (λ = 1.05) and additive (λ = 1.03) models were close to 1, indicating no inflation, thereby minimizing the risk of false-positive results. The identified variants were located in *TAS2R60* (in-frame deletion) and *SCNN1D* (one intronic and one missense variant), as shown in the volcano plot (**Figure 2B**). The intron variant in the *PLCB2* gene is located only 8 nucleotides away from the nearest exon, suggesting that it may play a role in splice site binding or be involved in gene regulatory mechanisms. Furthermore, variants in *SCNN1D* and PLCB2 exhibited mild linkage disequilibrium (0.2 < r^2^< 0.5) with other variants in their respective genes **(Figure S7**).

**Figure 2:**
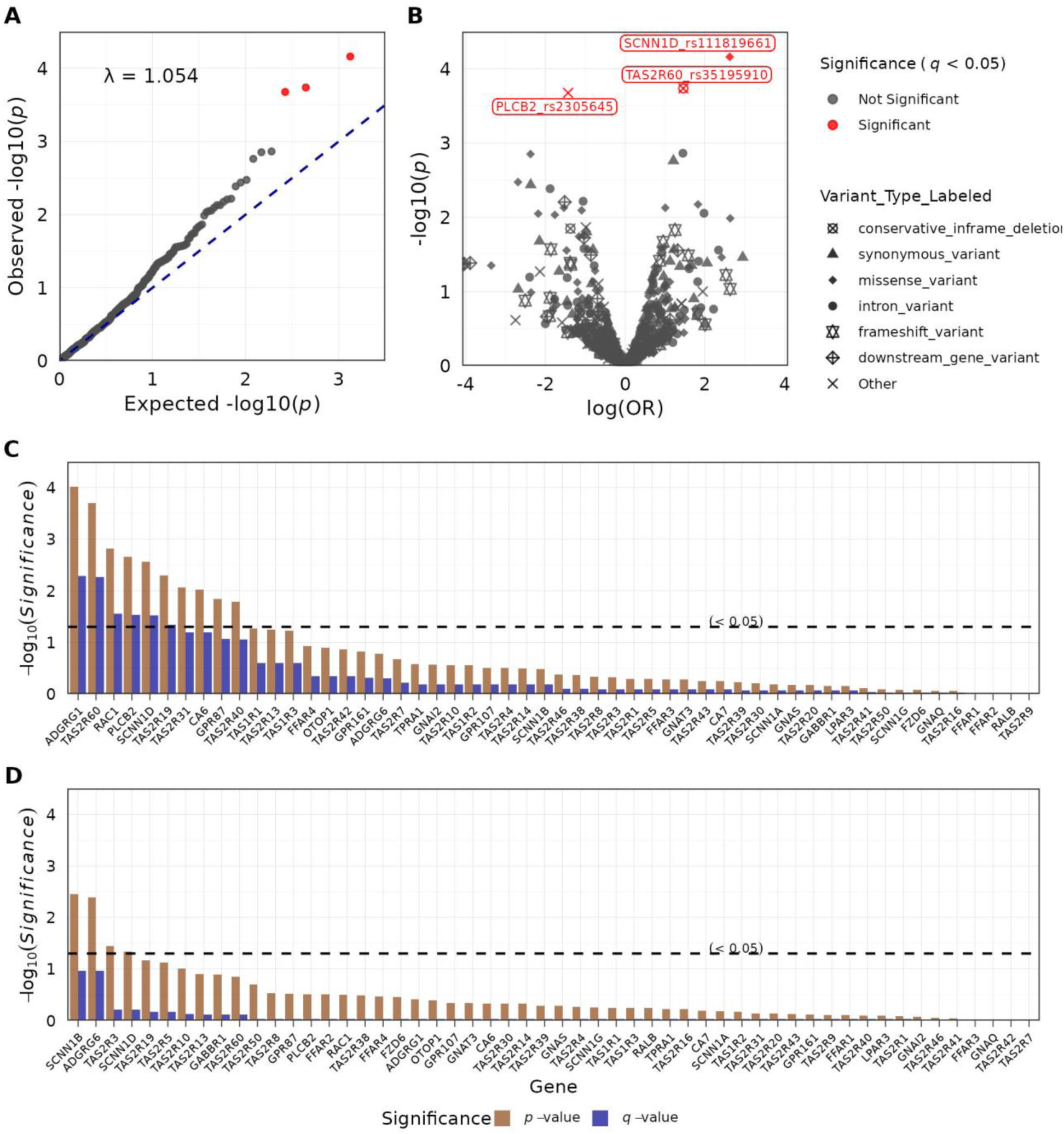
Taste genetic variants and genes association with ECC status. (A) The Q-Q plot illustrates the distribution of observed −log10(*p*-value) values against the expected −log10(*p*-value) values under the null hypothesis of no association between the variables. Each point represents a single nucleotide polymorphism (SNP). The blue dashed line indicates the null expectation, where the observed equals the expected. (B) Volcano plot. The x-axis represents the log2 odds ratio (OR), and the y-axis shows the −log10(*p*-value). Each point represents a variant, with the shape indicating the variant type. Red points denote significant associations for *q* < 0.05 (BH-adjusted *p*-value < 0.05). The Q-Q and volcano plots are from logistic regression models in PLINK under a dominant model for 55 taste-associated candidate genes, with age, sex, rural-urban status, SEFI score, and PCs 1–5 as covariates. (C) Gene-wise association for rare (minimum allele frequency, MAF < 0.01) and common (MAF > 0.01) variants with equal weights in the SKAT method in R. The bar plot shows *p*-values and *q*-values for the association between variants within each gene after removing singletons. The analysis was performed in the SKAT R-package using the SKAT-O option for optimal association, adjusting for age, sex, rural-urban status, SEFI score, and the top five principal components. The horizontal line marks the significance cutoff of 0.05. (D) Same as (C), but only for rare variants (MAF < 0.01).

**Table 2:** Significant loci identified in 55 taste-related genes to be associated with ECC status.

| **Dominant model** |  |  |  |  |  |  |  |  |  |  |
| --- | --- | --- | --- | --- | --- | --- | --- | --- | --- | --- |
| **Location** | **Gene** | **SNP_rsID** | **A1** | **A2** | **Frq_ECC** | **Frq_CF** | **OR** | ***p*** | ***p*_BH** | **Variant_Type** |
| chr1:1287578 | SCNN1D | rs111819661 | T | C | 0.175 | 0.020 | 6.165 | 6.91e-05 | 4.79e-02 | missense variant |
| chr7:143443837 | TAS2R60 | rs35195910 | G | GTCT | 0.397 | 0.179 | 2.749 | 1.83e-04 | 4.87e-02 | conservative<br>inframe deletion |
| chr15:40303364 | PLCB2 | rs2305645 | T | C | 0.151 | 0.295 | 0.371 | 2.11e-04 | 4.87e-02 | intron variant |
| **Additive model** |  |  |  |  |  |  |  |  |  |  |
| chr1:1287578 | SCNN1D | rs111819661 | T | C | 0.175 | 0.020 | 6.108 | 6.11e-05 | 4.24e-02 | missense variant |
| **Location**, Chromosomal location of the variant in the GRCh38 genome assembly.<br>**SNP_rsID**, Reference SNP ID number, if available.<br>**A1**, Allele corresponding to measured effect on the outcome.<br>**A2**, Reference allele, not corresponding to measured effect on the outcome.<br>**Frq_ECC/Frq_CF**, Allele frequencies in ECC and caries-free (CF) groups.<br>**OR**, Odds ratio, indicating the effect size.<br>***p***, *p*-value for the association.<br>***p*_BH**, Benjamini-Hochberg-corrected *p*-value for multiple testing.<br>**Variant_Type**, Functional annotation of the variant |  |  |  |  |  |  |  |  |  |  |

Sequence kernel association test (SKAT) analysis (S. Lee et al., 2012; Wu et al., 2011), which assessed gene-level associations by considering the cumulative effects of both common (minor allele frequency [MAF] >= 0.01) and rare (MAF < 0.01) variants, identified *ADGRG1* and *TAS2R60* as strongly associated with ECC outcomes at *q* < 0.05 (**Figure 2C**). The other significant genes were *RAC1*, *PLCB2*, *SCNN1D*, and *TAS2R19*. Although no significant genes were identified for rare variants after *p*-value adjustment, *ADGRG6* and *SCNN1B* showed suggestive associations before adjustment (**Figure 2D**).

### ECC microbiome and genetic variants

Microbiome associations for both bacteria and fungi with ECC were performed using Microbiome Multivariable Associations with Linear Models version 2(MaAsLin2) with a *q*-value cutoff of 0.01 (Mallick et al., 2021). After adjusting for age, sex, rural-urban status, and SEFI score, 39 bacterial species and two fungal species differed significantly between the CF and ECC groups (**Figure 3A**). At the genus level, ten bacterial genera and one fungal genus were identified as being significantly associated with ECC outcomes (**Figure 3B**).

**Figure 3:**
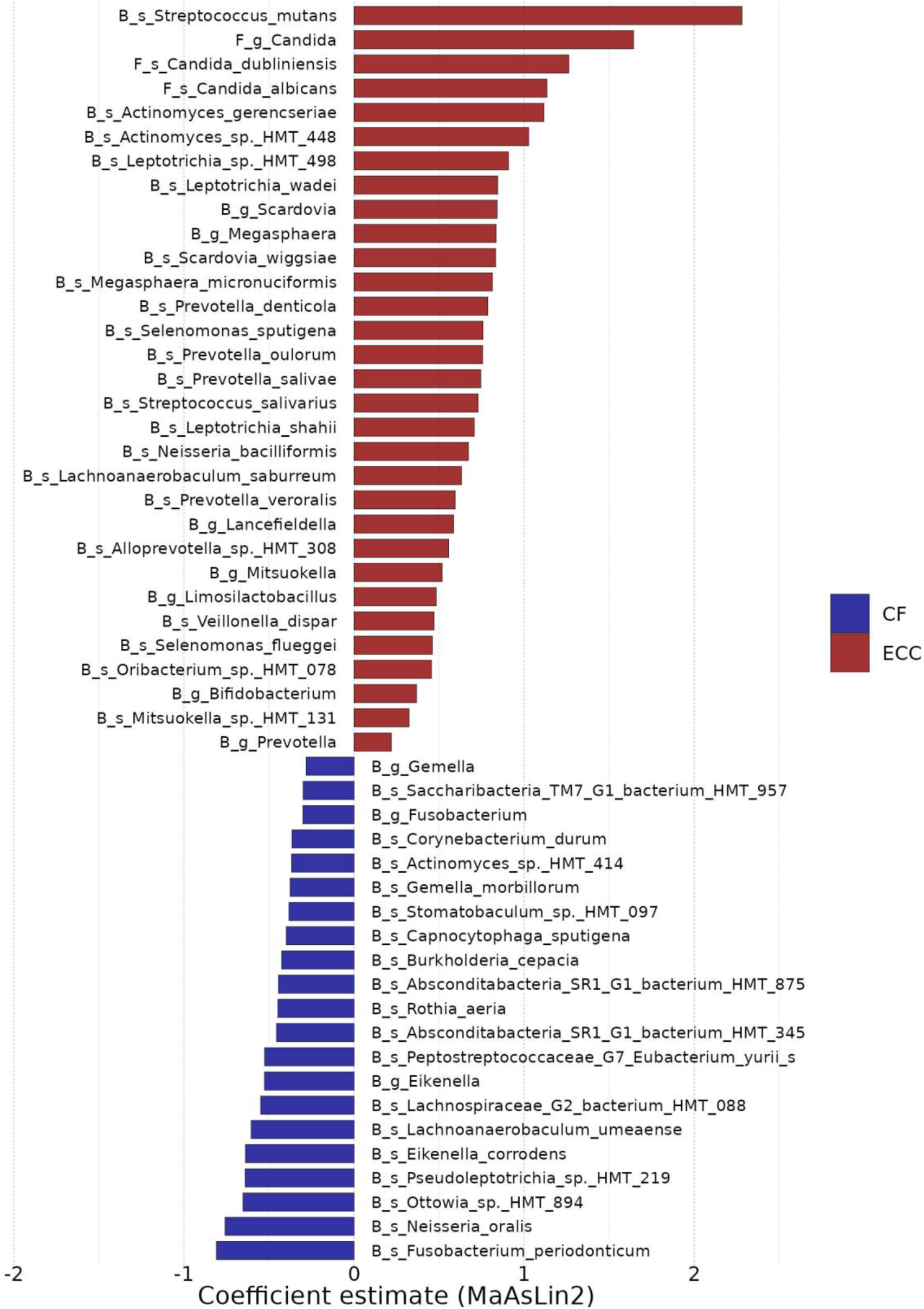
Association between dental plaque microbial taxa and ECC outcome. The coefficient estimates for significantly different taxa at *q*-values < 0.01 between the ECC and CF groups using the MaAsLin2 differential abundance analysis method after adjusting for age, sex, rural-urban status, and SEFI score. Microbial features are labeled as ‘B’ for bacterial and ‘F’ for fungal taxa. The prefix ‘s’ signifies species level and ‘g’ indicates genus level.

The most prominent bacterial associations included *Streptococcus mutans* and two *Actinomyces* species, *Actinomyces gerencseriae* and an unclassified species, *Actinomyces* sp.

HMT-448, which were strongly linked to the ECC condition. In contrast, *Fusobacterium periodonticum* and *Neisseria oralis* were significantly enriched in the CF samples. Among fungi, the genus *Candida* was the most abundant in ECC, although no specific fungal species were associated with CF conditions. Additionally, the bacterial genera *Scardovia* and *Fusobacterium* showed significantly increased abundance in ECC and CF groups, respectively.

### Microbiome quantitative trait loci analysis

Regarding the association between ECC-associated variants and dental plaque microbiome diversity, the three identified variants did not exhibit significant effects on bacterial alpha diversity (**Figure 4A**). However, variants in *SCNN1D* and *TAS2R60* were negatively correlated with the alpha diversity of fungal species, indicating an overabundance of certain fungal species in the presence of these variants. Additionally, these two variants were significantly associated with beta diversity in both bacterial and fungal datasets, whereas the PLCB2 variant was only associated with beta diversity in fungal samples (**Figure 4B**).

**Figure 4:**
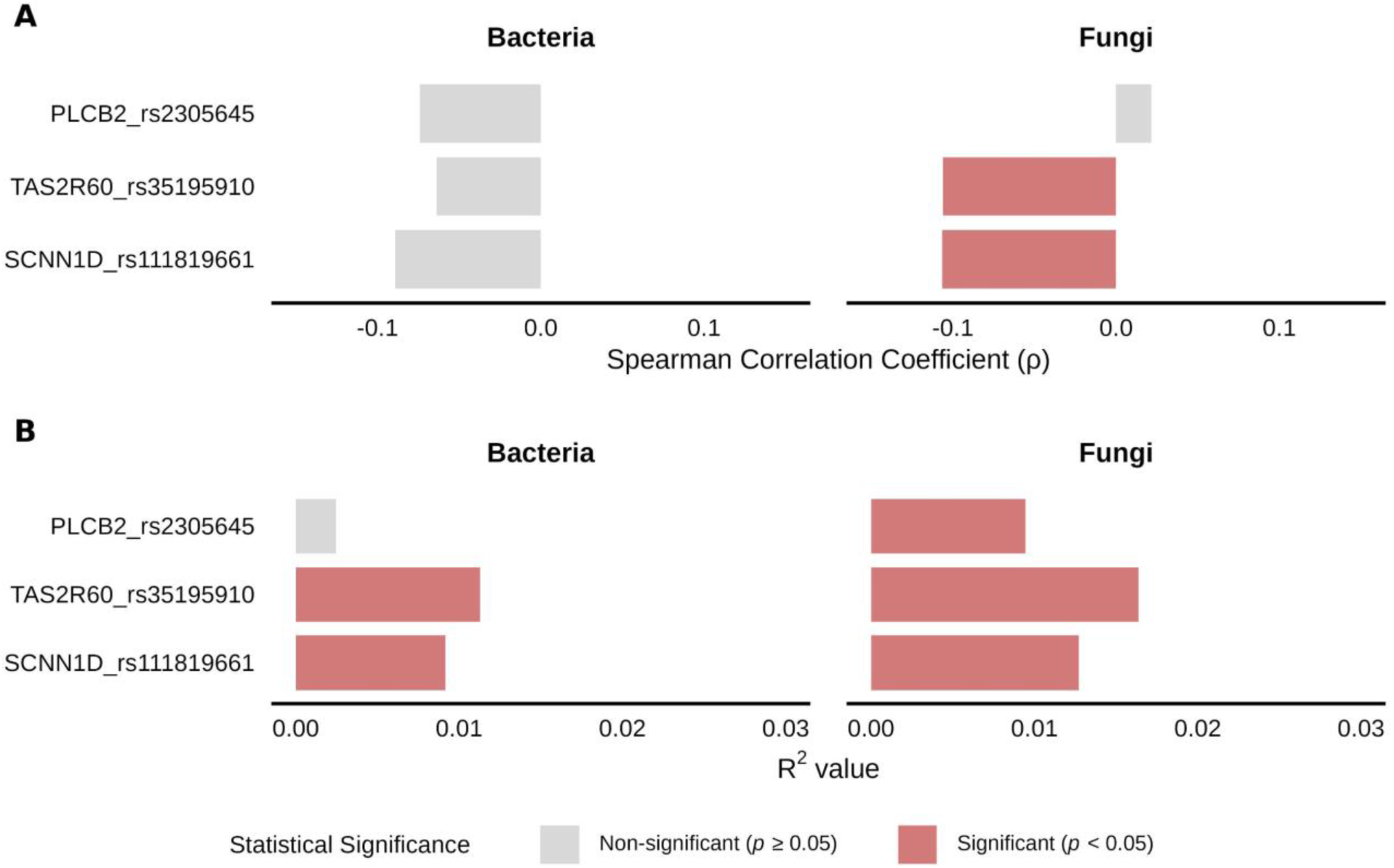
Spearman correlation of the ECC associated variants with alpha and beta diversity. (A) Significant genetic variants and their Spearman correlation coefficient (ρ) values for alpha diversity assessed using Shannon’s metric for bacteria and fungi. (B) Beta diversity analyses with significant genetic variants via PERMANOVA. The R^2^ values indicate the variance explained by the bacterial and fungal communities. The colored bar in each plot represents the statistical significance of the Spearman correlation at *p* < 0.05.

Microbiome quantitative trait loci (mbQTL) analysis was performed using linear associations with a dominant model for significant variants. At the genus level, no bacterial or fungal genera were found to be associated with any of the ECC-related variants (**Figure 5A**). The *TAS2R60*-rs35195910 variant was significantly associated with the bacterial species *Streptococcus intermedius*, *Parascardovia denticollens*, *Aggregatibacter* sp. HMT-898, and the fungal species *Protrudomyces lateralis* (**Figure 5B**). Meanwhile, the variant in *SCNN1D* was associated with one bacterial species, *Capnocytophaga* sp. HMT-878. However, none of these species were identified as being differentially abundant in ECC and CF.

**Figure 5:**
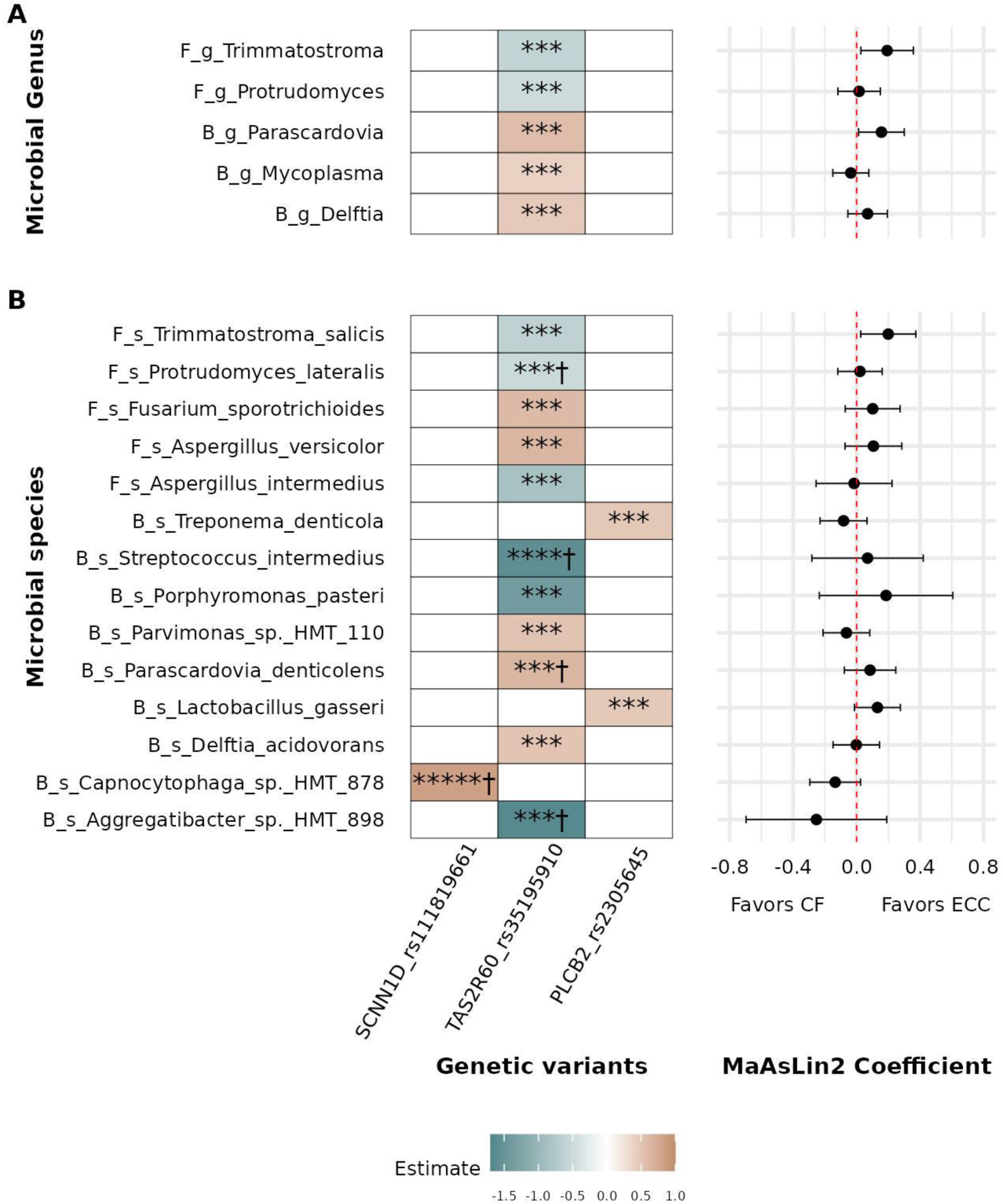
Association between ECC-associated variants and microbial taxa. The heatmap illustrates the association between ECC-associated genetic variants and microbial taxa when adjusted for covariates such as age, sex, urban-rural status, SEFI score, and the top five principal components from the genetic data. The forest plot shows the coefficient of association from the MaAsLin2 method between the selected microbial species in the heatmap and ECC outcome. Positive coefficients favor ECC, whereas negative coefficients favor CF status. The analysis was performed for significant variants identified in the dominant genetic model and all microbial species with CLR normalization using a linear model. The heatmap value signifies the estimates values, with red showing a positive association between variants with alternate alleles and green representing a negative association. The *p*-values are (* *p* < 0.05, ** *p* < 0.01, *** *p* < 0.001, **** *p* < 0.0001 ***** *p* < 0.00001), and “†“ indicates a significant association at BH adjusted *p* < 0.05. Microbial features are labeled as ‘B’ for bacterial and ‘F’ for fungal taxa. The prefix ‘s’ signifies species level and ‘g’ indicates genus level. (A) Genus level (B) Species level.

### Mediation analysis

The linear decomposition model (LDM) was used to test the potential mediation of multiple microbial species in the association between ECC outcomes and specific genetic variants (Yue & Hu, 2022). Three ECC-associated variants were examined to identify the mediating species from the differentially abundant species (**Figure 6A**). These results suggest that the association between the *TAS2R60* variant and ECC status may be mediated by four bacterial species: *Streptococcus mutans*, *Peptostreptococcaceae* subsp. *yurii margaretiae*, *Fusobacterium periodonticum*, and *Eikenella corrodens*. Additionally, the ECC outcome associated with the *SCNN1D* variant was mediated by *S. mutans*. Further analysis of the single variant with single-taxon mediation revealed only two significant mediators after adjusting for *p*-values: *S. mutans* and *Fusobacterium periodonticum*, both of which mediated the association between the *TAS2R60* variant and ECC (**Figure 6B**). However, the average mediation effect (ACME) of these two species was lower than the average total effect (ADE) of the variants on ECC outcomes, suggesting a partial mediation (**Figure 6C**).

**Figure 6:**
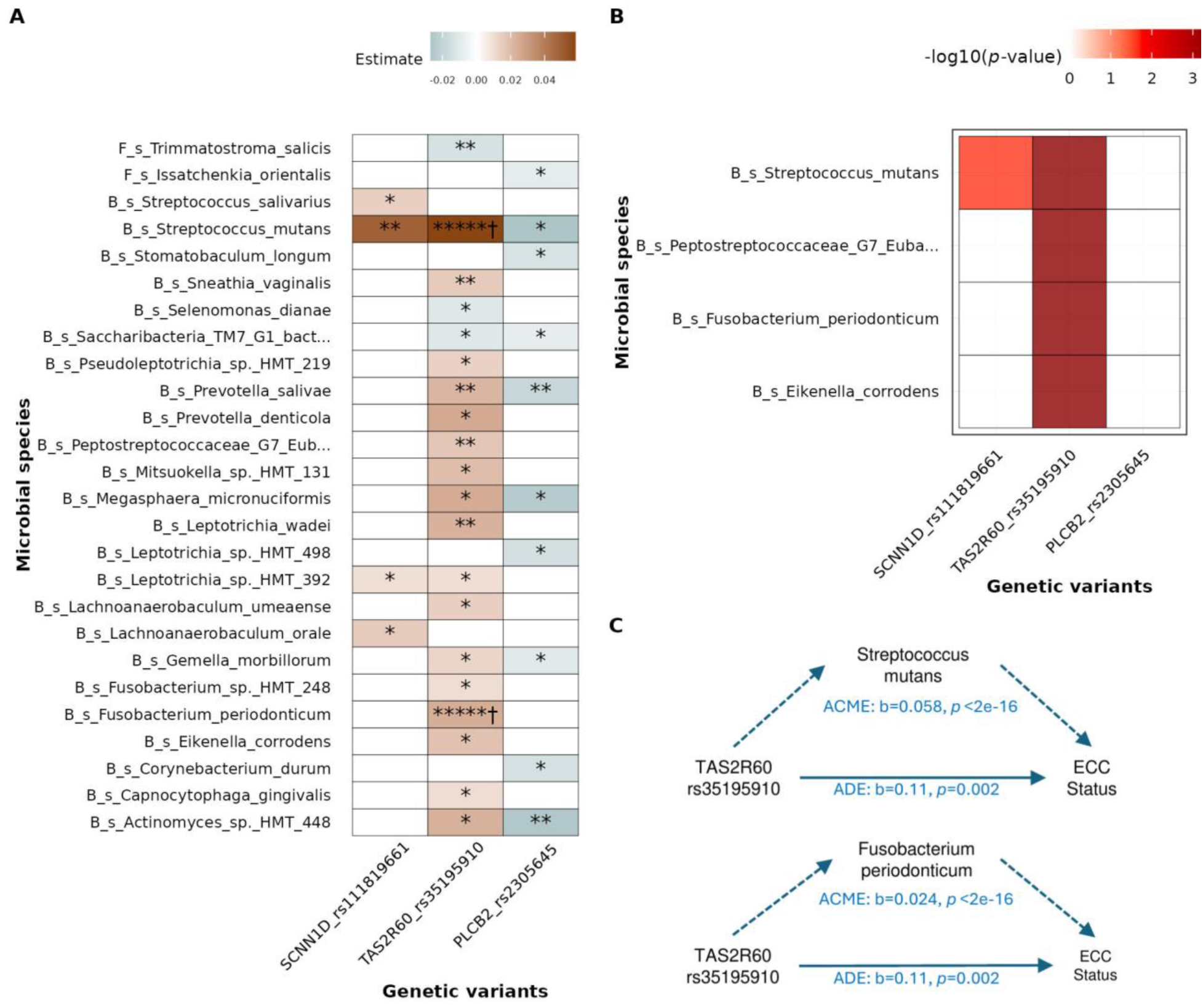
Mediation analysis for genetic variants and ECC association with microbial species as mediator. (A) Mediation analysis in the R package “mediation” for significant variants with each microbial species. The heatmap value signifies the estimate values, with red showing positive mediation between variants with alternate alleles and green representing a negative association. The *p*-values are (* *p* < 0.05, ** *p* < 0.01, ***** *p* < 0.00001), and “†” indicates a significant association at BH adjusted *p*<0.05. (B) Mediation was performed using LDM with one SNP and multiple mediator approach. This heatmap illustrates the significant mediation of microbial species between genetic variants and ECC outcomes. The x-axis shows the genetic variants, and the y-axis lists the microbial species. The color scale represents the negative log10 of *p*-values. (C) Average mediation effect (ACME) and average direct effect (ADE) for significant mediators at BH adjusted *p*<0.05 in (A). Microbial features are labeled as ‘B’ for bacterial and ‘F’ for fungal taxa. The prefix ‘s’ signifies species level and ‘g’ indicates genus level.

### ECC prediction models

To assess the predictability of ECC outcomes, four machine learning (ML) methods were trained and tested on each dataset (microbiome, genetic variants, and covariates) individually and in combination. Among the methods, the Random Forest (RF) model demonstrated comparable or superior performance compared to the other models (**Figure 7A**). Using RF, the prediction of ECC outcomes with microbiome data achieved the best classification performance, with the Area Under the Receiver Operating Characteristic curve (AUROC) and Area Under the Precision-Recall Curve (AUPRC) values both reaching nearly 0.94. For the genetic data, RF provided AUROC and AUPRC values of 0.85 and 0.89, respectively. The covariate-based model, which included age, sex, rural-urban status, and SEFI score, yielded an AUROC of 0.8.

**Figure 7:**
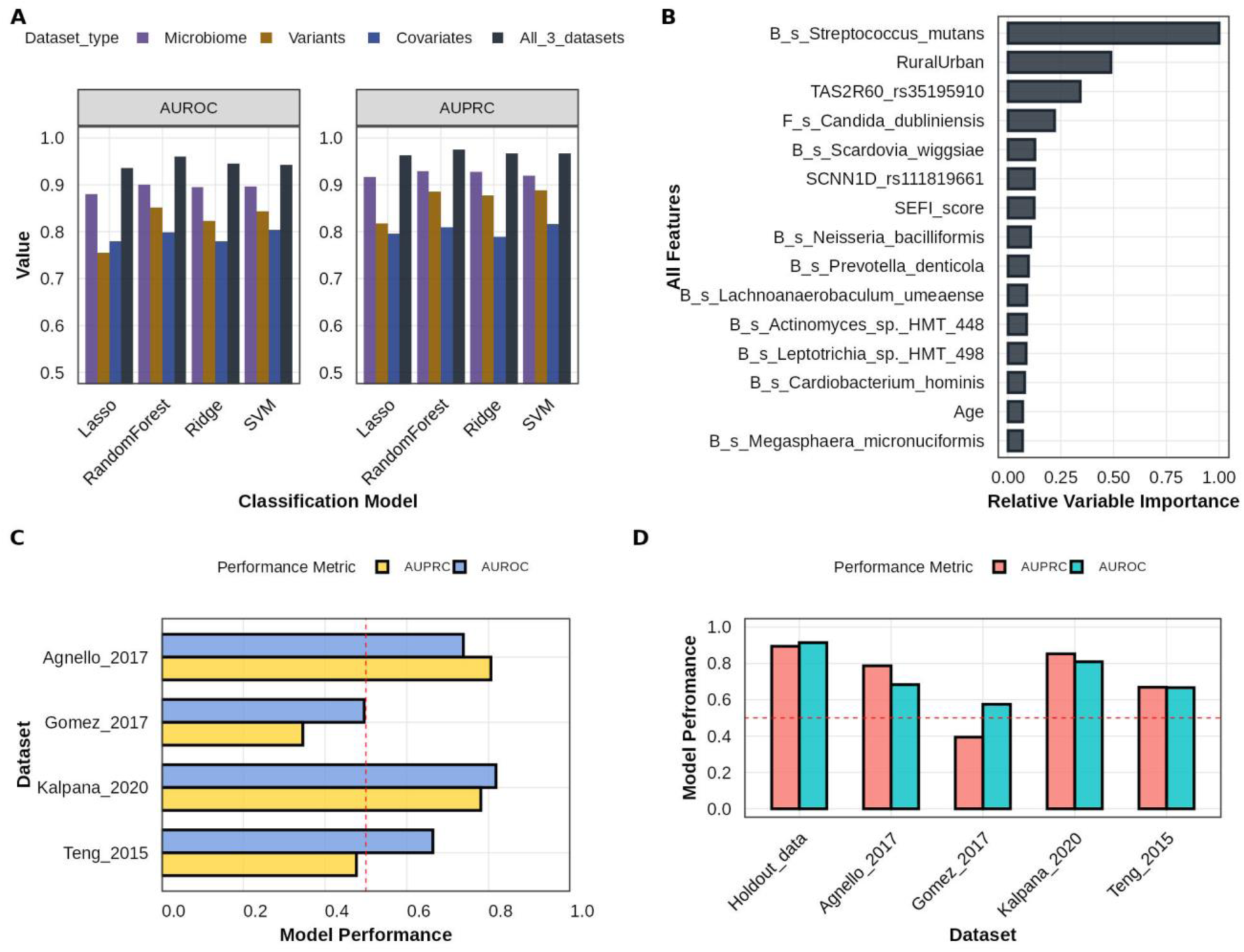
Machine learning based classification for ECC status using microbiome, genetic variants and covariate data. (A)Comparison of machine learning models based on individual and combined datasets for microbiome (fungi and bacteria), genetic variants, and covariate data using AUROC and AUPRC metrics. The ML models selected for analysis were lasso, random forest (RF), ridge, and SVM. (B) The RF model with the combined dataset was chosen to plot the variable importance bar plot. The bars indicate the relative importance of each feature when all features are used to classify ECC_status. (C) RF model performance on sPLSDA batch-corrected external datasets. Four external datasets were chosen to assess the RF model based on only microbiome data, and the model performance was assessed using AUROC and AUPRC values. (D) ECC prediction model performance on sPLSDA batch-corrected external datasets based on the MRS score on CLR-normalized datasets. Microbial features are labeled as ‘B’ for bacterial and ‘F’ for fungal taxa. The prefix ‘s’ signifies species level and ‘g’ indicates genus level.

When features from the microbiome, genetic variants, and covariates for the prediction of ECC status were combined, the model performance for the prediction of ECC status for the prediction of ECC status improved to 0.96 for both AUROC and AUPRC. Feature importance analysis in the RF model identified the top predictive features from all three data sets. The most influential factor was the bacterial species *S. mutans*, followed by urban-rural status and the variant *TAS2R60*-rs35195910 (**Figure 7B**). The fungal species *Candida dubliniensis* ranked among the top five variables in the combined dataset and was the second most important feature in the microbiome data after *S. mutans*. Among the other covariates, age ranked in the top variables, whereas sex was not identified as a significant feature.

The microbiome RF model was further tested using external microbiome datasets with sPLSDA batch correction (**Figure 7C**) (Agnello et al., 2017; Gomez et al., 2017; Kalpana et al., 2020; Teng et al., 2015). The AUROC values for the test performance were 0.78, 0.82, and 0.71 for the Agnello-2017, Kalpana-2020, and Teng-2015 datasets, respectively. However, the Gomez-2017 dataset exhibited near-random performance with an AUROC of 0.47. For AUPRC, the performances of the Gomez-2017 and Teng-2015 datasets were lower than their respective AUROC values. Using sPLSDA with batch-corrected data, the performance slightly improved for the Kalpana-2020 dataset compared to the CLR-normalized values (**Figure 7C and S11A**).

However, a decrease in AUPRC performance was observed for the Teng-2015 dataset, which decreased from 0.54 to 0.48.

The predictive performance of the microbial risk score (MRS) was evaluated using batch-corrected datasets. On the holdout data from this study, the MRS demonstrated a strong performance, achieving AUROC and AUPRC values close to 0.9 (**Figure 7D**). The performance remained high on the Kalpana-2020 dataset, with an AUROC exceeding 0.8, and was moderate on the Agnello-2017 dataset, with an AUROC of approximately 0.75. In contrast, the performance was lower on the remaining datasets, with AUROC values below 0.68 and 0.58 for Teng-2015 and Gomez-2017, respectively. MRS was also tested on non-batch-corrected datasets (Figure S11B). While the AUROC values remained largely unchanged compared to the non-batch-corrected data, the AUPRC values showed slight improvements (**Figure 7D**). This was particularly evident in the two datasets with lower AUROC values: Teng-2015 and Gomez-2017 datasets.

## Discussion

This study examined the relationship between taste genetics and the oral microbiome in ECC in the context of sociodemographic factors using an integrative analysis. We focused on taste-associated genes because of their impact on taste preferences, which influence dietary habits and subsequently shape the oral microbiome. Previous research has suggested potential interactions between receptors encoded by these genes and genetic variants that could modify these interactions. This study employed robust statistical analyses, incorporating age, sex, SEFI score, and rural-urban status as covariates, to refine the identification of potential genetic variants and their association with the dental plaque microbiome. Our findings identified genetic variants in the bitter taste receptor gene *TAS2R60* (rs35195910), sodium ion transport gene *SCNN1D* (rs111819661), and phospholipase gene *PLCB2* (rs2305645) that were associated with ECC. These results align with findings from large population studies that highlight the role of host taste genetics in ECC (Orlova et al., 2022; Shrestha et al., 2024). Furthermore, these variants were linked to changes in the microbial species within the dental plaque microbiome of the participants.

Focusing on specific genes allows us to sequence them using deep sequencing methods, which is more logistically feasible than genome-wide analysis. Deep sequencing facilitates the identification of novel variants, particularly frameshifts caused by insertions and deletions. Some novel variants, such as those in *TAS2R19* and *TAS2R42*, appeared significant in the base model adjusted for age and sex but lost significance when adjusted for additional covariates. Notably, the single nucleotide variation in *SCNN1D*-rs111819661 demonstrated both additive and dominant roles (**Table 2**). This dual mode of association has also been reported in genetic studies on rheumatoid arthritis and cognitive function in cancer patients treated with opioids (Jiang et al., 2012; Kurita et al., 2016). However, larger sample sizes are required to empirically determine whether this model reflects an additive or a dominant effect.

Additionally, we analyzed the cumulative association of both rare and common variants and found that variants in *ADGRG1* and *TAS2R60* were associated with ECC. The association with ECC may be attributed to the highly ECC-associated variant rs35195910 in *TAS2R60*, whereas the significance of *ADGRG1* highlights the potential polygenic nature of this relationship.

However, none of the genes showed a significant association with ECC when only rare variant analyses were performed (**Figure 2D**). This finding suggests that rare variants, even in combination, may not influence ECC outcomes and that the association with *ADGRG1* is likely driven by multiple common alleles acting in combination.

The role of host genetics in ECC has yielded varying results in previous studies. While some studies suggest a potential role for host genetic composition, others have found minimal or no influence, emphasizing the dominant roles of environmental factors and time in shaping the microbial composition (Freire et al., 2020; Gomez et al., 2017; Mukherjee et al., 2021). A recent meta-analysis of genome-wide association studies (GWAS) and transcriptome-wide association studies (TWAS) for ECC did not identify associations with variants in taste-associated genes.

However, their TWAS analysis highlighted the association between the expression of several bitter taste receptors and ECC (Orlova et al., 2022). Additionally, the pathway related to taste receptor activity was found to be associated with ECC, and the GWAS association of the variant rs58016156 in *KCNU1*, which is involved in sweet taste signaling, further underscores the role of taste genetics (Shrestha et al., 2024).

According to the Genome Aggregation Database (gnomAD) (Karczewski et al., 2020), the frequencies of the significant variants *SCNN1D-*rs111819661 and *PLCB2-*rs2305645 in our samples were within the range of the average allele frequencies observed in the general population. In contrast, the variant rs35195910 in *TAS2R60* demonstrated a higher frequency in our CF samples and an even greater prevalence in ECC samples, exceeding the population average (**Figure S8**). This suggests that the population in consideration may be at a higher risk of rs35195910 associated ECC outcomes.

We did not identify any ECC-associated variants that showed a significant relationship with ECC-causing microbial species in the linear association analysis, including ECC status as one of the covariates. Although some dental plaque species were significantly associated with ECC-related variants, none of the taxa at the genus level exhibited significant associations. This suggests that interactions between receptors and microbes may occur at the species level, rather than at the genus level. Both mbQTL and mediation analyses indicated that *TAS2R60* influences a greater number of species than the other two variants (**Figure 5A, 6B**). *TAS2R60* is an orphan taste receptor with no known agonists or antagonists. The high prevalence of this variant and its association with ECC and ECC-related microbiota highlight the importance of future research on the role of this receptor in host physiology and microbial interactions (Kouakou & Lee, 2023; Medapati, Singh, et al., 2021; Xi et al., 2022). In the stratified analysis of the ECC and CF samples, only one association, between *TAS2R60-*rs35195910 and *Streptococcus intermedius*, was common in both groups (**Figure S9**). Furthermore, the mediation observed between *S. mutans* and the bitter taste receptors, T2Rs, may be through competence-stimulating peptides secreted by *S. mutans* (Medapati, Singh, et al., 2021). As previously identified, competence stimulating peptide-1 interaction with the bitter taste receptor induces an immunogenic response, such as the secretion of CXCL-8/IL-8, TNF-α, and IL-6, and the inhibition of other oral microbes (Medapati, Bhagirath, et al., 2021; Medapati, Singh, et al., 2021).

The top ECC-associated species belong to genera such as *Streptococcus*, *Actinomyces*, and *Candida*, which have been previously linked to biofilm formation (Karkowska-Kuleta et al., 2022; Rickard et al., 2003). A recent study demonstrated associations between *Staphylococcus epidermidis* and the host genetic variation rs3828054 in the *TUFT1* gene, as well as between Candida species and the enamel gene *ENAM* variant rs3796704, suggesting a connection between host genes and microorganisms (Vieira & Modesto, 2022). In our study, despite the strong association between *Candida* species and ECC, no taste gene variants were found to be associated with *Candida* species. However, the significant negative correlation between both *TAS2R60*-rs35195910 and *SCNN1D-*rs111819661 with fungal diversity suggests the enrichment of certain species, potentially *C. dubliniensis* and *C. albicans*. Furthermore, although not statistically significant, we observed a negative association between the CF-associated species *Fusobacterium periodonticum* and a novel variant in *TAS1R1* at position 6574730 (**Figure S10**). This finding suggests that the presence of this variant reduces the abundance of ECC-protective species in the dental plaque microbiome.

Our results align with those of Gomez et al., where some microorganisms were associated with host genetics, but cariogenic species were not (Gomez et al., 2017). Additionally, our previous research demonstrated that many of these taxa are influenced by socioeconomic and environmental factors, such as the SEFI score, rural-urban status, and bedtime snacking habits (M. W. Khan, de Jesus, et al., 2024). Nonetheless, certain ECC-associated species may partially mediate the relationship between host genetics and disease outcomes.

Machine learning models enable the integration of different data types, such as microbiome, genetic, and covariate data, into a single model. Additionally, models such as RF can capture the nonlinear patterns and variable interactions in the data. By investigating these models, we can assess the relative importance of each variable while controlling for others. The ranking of *S. mutans* as the top feature underscores its critical role in ECC etiology (**Figure 7B**). The appearance of covariates, specifically the SEFI score among the top-ranked variables, highlights their association and the need to adjust for these factors when studying ECC. Although previous studies have highlighted the role of SES, our study compares these factors with microbial and taste genetic factors, providing the relative importance of these variables through an integrative model.

Our RF model demonstrated that genetic variants alone can predict ECC outcomes, with an AUROC of 0.85 (**Figure 7A**). Although this is lower than the predictive performance achieved using microbiome data, the appearance of genetic variants, such as *TAS2R60* and *SCNN1D*, among the top features suggests that combining genetic and microbiome data can improve microbiome-based risk prediction. This finding also indicates that certain genetic markers may play a more significant role than some ECC-associated microbial species in shaping the etiology of ECC, further supporting the influence of taste genetics on caries progression. This is further evident from a Polish study that showed an AUC of 0.97 for ECC prediction using candidate genes (Zaorska et al., 2021). Our results add to the potential candidate genetic markers that may further improve ECC prediction.

Moreover, MRS, which is based on a linear combination of ECC-associated microbial markers, achieved comparable performance on both the internal and external datasets. These results validate the robustness of microbial markers in ECC risk assessment. A combined ECC risk score, incorporating both microbial and genetic markers, could potentially improve predictive accuracy in external datasets where both genetic and microbiome data are available.

Furthermore, as suggested by Blostein et al., MRS can be used as one of the covariate to identify the polygenic burden of ECC-associated variants (Blostein et al., 2023). However, given the observed differences in the microbiome and genetic associations with ECC, the generalizability of these methods must be evaluated across diverse populations.

Future multiomics approaches, such as combining metatranscriptomics and metabolomics for both host and microbial communities, could provide deeper insights by capturing differences in gene expression and highlighting the effects of genetic variants on host gene expression and microbial abundance (Divaris et al., 2019). These interactions may also extend to adult caries, as some genetic markers are consistent between ECC and caries experience in adulthood (Shrestha et al., 2024). Furthermore, integrating gene-environment studies, such as those on host-specific dietary influences, may further elucidate the environmental factors contributing to ECC. Collecting and including severity scores for ECC and incorporating standardized subtyping based on calibrated dentists’ scoring systems is also suggested (Gormley et al., 2023; Simancas-Pallares et al., 2023). Moreover, recent advancements in deep learning methods for receptor-ligand interactions could facilitate the development of algorithms to study microbial ligands and their interactions with taste receptors. These interactions can be validated through laboratory testing, offering potential insights into the additional mechanisms underlying ECC.

This study may have some potential limitations. As the study design is cross-sectional and not longitudinal, it will be difficult to determine whether the alteration of the microbiome is the cause or consequence of the disease. The external validation of our machine learning models showed moderate performance, suggesting that microbial profiles may not generalize well across different geographic populations owing to variability in microbial composition. Moreover, the limited association between the identified variants and ECC-associated microbes does not eliminate the possibility of interactions or the influence of these variants on such associations.

Since our analyses relied on linear associations between variants and microbial abundance, which may not fully reflect biological complexity, further research using host transcriptomics and microbial metatranscriptomics is required (Chetty & Blekhman, 2024).

## Conclusion

While the role of the microbiome has been extensively studied in previous research, our study extends the knowledge of the genetic basis of ECC by focusing on taste genes along with socioeconomic variables. Genetic factors may influence ECC through host protein interactions with specific microbial species, potentially leading to microbial dysbiosis. Although ECC outcomes are primarily determined by the oral microbiome, certain genetic variants rank highly as biomarkers in our multiomics models. Our findings provide a rationale for integrated studies using both microbiome and host genetics, while considering socioeconomic conditions as covariates to study ECC etiology. These results can help identify susceptible individuals, enabling personalized therapeutic approaches and informing dental public health policymakers and oral healthcare providers to implement targeted early interventions at both societal and familial levels.

## Methodology

### Experimental model and study participant details

This study included 538 participants under the age of 72 months, recruited between December 2017 and July 2022 from the Children’s Hospital Research Institute of Manitoba (CHRIM), Misericordia Health Center (MHC), and various community clinics across Winnipeg, Manitoba. The eligibility criteria required participants to be less than 72 months of age and not currently on antibiotics. In this cross-sectional study, samples from children with ECC were considered cases, whereas those from children with CF were considered controls. ECC was diagnosed by experienced dentists following the American Academy of Pediatric Dentistry (AAPD) definition, which identifies ECC as the presence of one or more decayed, missing (due to caries), or filled tooth surfaces in any primary tooth (AAPD, 2024). A questionnaire was completed by parents or caregivers gathering additional information, including the child’s age, sex, place of residence (urban or rural), feeding habits, and mode of birth. The socioeconomic factor index (SEFI), an area-based socioeconomic measure, was derived from the Manitoba Centre for Health Policy (MCHP) data based on participants’ postal codes (S. Lee et al., 2012). SEFI is derived from Canadian Census data and is a combined score of four areas: unemployment rate, average household income, proportion of single-parent households, and proportion of the population without a high school graduation. When interpreting the SEFI, a lower score indicates a more favorable SES, while a score greater than 0 reflects less ideal socioeconomic conditions.

### Sample collection, DNA extraction, and sequencing

Samples were collected from two sites for each participant: supragingival plaque (dental plaque) and buccal swabs to sequence the microbiome and host genetic factors, respectively. Dental plaque samples were obtained using sterile interdental brushes and collected from all tooth surfaces, regardless of cavity presence (Agnello et al., 2017; de Jesus et al., 2020; M. W. Khan, de Jesus, et al., 2024). Buccal cavity samples were collected by swabbing the buccal mucosa and the anterior floor of the mouth beneath the tongue (de Jesus, Khan, et al., 2021).

Amplicon sequencing of bacteria and fungi was performed using MiSeq or NovoSeq6000 platforms (Illumina Inc., San Diego, CA, USA). For bacterial 16S rRNA (V4 region), the primer pair 515F (5′-GTGCCAGCMGCCGCGGTAA-3′) and 806R (5′-GGACTACHVGGGTWTCTAAT-3′) was used. Fungal ITS1 regions were targeted using the primers ITS1-30 (5′-GTCCCTGCCCTTTGTACACA-3′) and ITS1-217 (5′-TTTCGCTGCGTTCTTCATCG-3′). Raw sequences were processed using QIIME2 to generate an ASV table containing microbial abundance data.

For host genetic analysis, 55 candidate genes were selected based on their roles in taste signal transduction, including receptors for bitter, sweet, umami, salt, sour, carbonation, and fatty acids, and their associated signaling proteins (**Table S1**). DNA was isolated using the QIAamp Mini Kit (Qiagen, Hilden, Germany) according to the manufacturer’s protocol. The candidate genes were amplified using primers designed for the Fluidigm platform to facilitate Illumina sequencing, ensuring that each amplified region covered approximately 250–300 base pairs for compatibility with 150 × 2 paired-end sequencing. All sequencing was conducted at the Genome Quebec Innovation Center (Montreal, Canada), and the resulting sequences were provided as demultiplexed paired-end FASTQ files with the barcodes removed. Additional details regarding these methods are given the our previous publications (de Jesus, Khan, et al., 2021; de Jesus et al., 2022)

### Raw sequence processing

#### Microbiome

The raw FASTQ sequences from amplicon sequencing were processed separately for 16S rRNA and ITS1 in QIIME2 to generate amplicon sequence variant (ASV) tables for bacteria and fungi (M. W. Khan, Fung, et al., 2024). For bacterial analysis, the DADA2 plugin in QIIME2 was used, whereas fungal analysis included an additional preprocessing step using Q2-ITSxpress in QIIME2 to remove conserved regions flanking ITS1 before applying the DADA2. Taxonomic assignment was conducted using the Human Oral Microbiome Database (HOMD, version 15.23) for bacteria and UNITE (version 9.0) for fungi. Taxonomies with less than 5% prevalence across the samples were excluded from the final ASV tables.

Differentially abundant species were identified using the MaAsLin2 method, which performs multivariable associations while accounting for confounding variables (Mallick et al., 2021). This analysis used ECC status as the outcome variable, with age, sex, rural-urban status, and SEFI score as the confounders. MaAsLin2 applies centered log-ratio (CLR) normalization internally before performing association tests and provides coefficients along with *q*-values (Gloor et al., 2017), which are *p*-values adjusted using the Benjamini-Hochberg (BH) method. In this study, MaAsLin2 was applied separately to the bacterial and fungal ASV tables with a significance threshold of *q* < 0.01. A detailed description of microbiome analysis using amplicon sequencing and preprocessing steps to obtain taxonomic abundances at the species and genus levels has been provided in our recent publication (M. W. Khan, de Jesus, et al., 2024).

#### Candidate genes

The raw sequences of the candidate genes were mapped to the GRCh38 human reference genome using the BWA mapping tool (Li & Durbin, 2009). The workflow adhered to the GATK best practice guidelines, with the exception of PCR duplicate removal, as this step is not recommended for targeted region amplification (Ebbert et al., 2016; Fu et al., 2018; McKenna et al., 2010). A schematic of the GATK pipeline steps, including the integration of data from multiple batches during the analysis, is provided in the Supplementary File (**Figure S1A**).

A combined variant calling format (VCF) file containing genotyping information for the genes included in the study was generated for all 538 samples. This file was cleaned and filtered using VCFtools (v 0.1.16). Quality control steps included filtering for a Phred quality score greater than 25, missing data less than 50%, and a sequencing depth greater than 5. SNPs were annotated using BCFtools with the human dbSNP138 database, and variant annotation was performed using SnpEff based on the human genome version, GRCh38.

For missing variants, all missing values were considered as not available. For analyses in which missing genotypes were not permissible, the genotype of the reference genome was used.

Genetic variants were named using the Gene_rsID approach when the rsID was available. For variants lacking rsID, the naming convention Gene_p followed by the chromosomal position was used. The absence of rsID also indicates novel variants not assigned to the dbSNP138 database.

#### Genetic association testing

The association between the variants identified by the GATK pipeline and ECC was tested using PLINK (v1.90) (McKenna et al., 2010; Purcell et al., 2007). Quality control and variant selection criteria included a genotype missingness threshold of less than 1%, a minor allele frequency (MAF) of at least 1%, and the removal of SNPs that significantly deviated from the Hardy-Weinberg equilibrium (HWE) with a *p*-value threshold of 0.00001. A logistic regression approach in PLINK was used to assess the association between genetic variants and ECC status.

The analysis was performed using both additive and dominant genetic models, wherein the genotypes were coded as 0, 1, and 2 in the additive model and 0 and 1 in the dominant model. To identify the appropriate covariates, a base model was first constructed in PLINK, including only age and sex as covariates. Additional covariates, such as rural-urban status, SEFI score, the first five PCs, and batches, were subsequently added to compare their AIC and BIC values using a linear model. The model was tested using the top five variants identified in the base model and the genomic inflation factor, which quantifies the deviation of the observed *p*-values from those expected under the null hypothesis. The AIC and BIC criteria for model performance for covariate combinations suggested that while rural-urban status and SEFI score improved the model, batch information did not (**Figure S4**). The final model included age, sex, urban-rural status, SEFI score, and top five PCs. Logistic regression in PLINK was performed with a *q*-value (Benjamini-Hochberg adjusted *p*-value) below 0.05 considered significant.

Additionally, gene-level association analyses of ECC outcomes were conducted using the SKAT method. Equal weights were applied to all variants, including those with MAF greater than or less than 0.01. However, for gene-level analyses focusing only on rare variants, an MAF threshold of less than 0.01 was used. The SKAT analysis employed the “SKAT-O” option, which optimally combines the standard SKAT test and burden test, adapting to variant causality and effect directionality (S. Lee et al., 2012; Wu et al., 2011).

#### Microbiome Quantitative Trait Loci Analysis

We first examined the changes in microbial diversity at both the alpha and beta levels associated with the three variants identified as ECC-associated. Microbial alpha diversity was measured using the Shannon index in Phyloseq, and correlations between Shannon diversity and variants were assessed using Spearman’s correlation (**Figure 4A**). For beta diversity, the distance matrix was calculated using the Bray method in Phyloseq, and the variance in the distance matrix explained by genetic variants was analyzed using the “adonis2” function in Vegan.

To perform the microbiome quantitative trait loci (mbQTL) analysis, linear regression analysis was performed using the “glm” function. In this analysis, variants were treated as independent variables, and microbial abundance served as the outcome variable, with ECC status included as an additional covariate along with the other covariates used in the genetic association testing.

The inclusion of ECC status as a covariate accounted for its potential confounding effect on microbial abundance, thereby reducing bias in the associations. The *p*-values were adjusted using the BH method. Only significant variants identified under the dominant genetic model were selected for analysis. To further explore these associations, a stratified association analysis was conducted separately within the ECC and CF samples using the same methodology.

## Mediation Analysis

To determine whether the effect of genetic variants on ECC outcomes was mediated through the microbiome, we used LDM mediation in R, which applies a linear decomposition model to test the mediation effects (Yue & Hu, 2022). This approach evaluates the exposure-taxon and taxon-outcome associations, where the exposure is an individual genetic variant and the outcome is ECC status. This analysis assumes that each variant independently influences several microbial species. Therefore, while the microbiome data included all significant taxa identified using MaAsLin2, only one variant was analyzed at a time.

LDM mediation enables mediation testing at the community level, incorporating all taxa provided in the analysis. The results identified potential taxa mediating the relationship between a genetic variant and ECC outcomes based on a defined significance cutoff. All significant taxa identified using the MaAsLin2 method were tested for their mediating effects. Furthermore, we conducted mediation analysis using the R package mediation to evaluate the mediation effect between a single genetic variant and a single microbial species. This approach provided the average causal mediation effect (ACME) as the mediation coefficient and the average direct effect (ADE) to quantify the direct relationship.

### Machine Learning Classifiers for ECC

To evaluate the relative importance of genotyping variants identified in our study alongside microbiome features, we incorporated genotyping data for quality filtered variants in a dominant fashion with microbiome data and constructed a classifier for CF and ECC classification in R using the tidymodels package (version 1.1.1). The RF, Lasso, Ridge, and SVM models were tested owing to their frequent application to similar data types. The model was tuned for the minimum node size, number of trees, and number of variables at each split. A grid size of 50 was used for the hyperparameter tuning.

The model performance was assessed using the AUROC and AUPRC metrics. The AUROC value quantifies the models’ ability to correctly predict the outcome classes, whereas the AUPRC suggests a trade-off between the precision and recall of the prediction. The variable importance in the final RF model was evaluated based on the mean decrease in accuracy upon shuffling each feature using the VIP package (version 0.4.1) in R to measure the impact on performance. Additionally, the RF model trained on bacterial species from our data was tested on external bacterial datasets from our previous study (M. W. Khan, Fung, et al., 2024), and the performance was assessed using the same criteria as above. The prediction was conducted with and without batch correction to assess performance under different preprocessing conditions.

### Microbiome Risk Score

In addition to the machine learning methods described in the previous section, we utilized MRS markers based on bacterial species that can be tested on external datasets. Similar to the polygenic risk score (Choi et al., 2020), the MRS score calculates an individual’s predisposition to ECC based on their microbiome profile by aggregating the effects of multiple microbial species identified as ECC-associated by MaAsLin2, with a *q*-value cutoff of < 0.01 (Choi et al., 2020). Each microbial marker was weighted by its MaAsLin2 coefficient and combined in a linear equation with the microbial profile. The MRS was calculated for 80% of the samples used in our study, and the resulting score represented the cumulative risk. The equation was then used as a predictor for the remaining 20% of the holdout test dataset. The performance of the MRS derived from our dataset was evaluated using four external datasets. Predictions using MRS were assessed using the AUROC and AUPRC scores.

## Acknowledgments

This study was funded by the Canadian Institutes of Health Research (CIHR) through an operating grant PJT-159731. P. H. is supported by the Canada Research Chairs Tier II Program (CRC-2021-00482). R. J. S. holds a CIHR Applied Public Health Chair in “Public Health Approaches to Improve Access to Oral Health Care and Oral Health Status for Young Children in Canada”. Both M.W.K. and V.C.J. received GETS funding from the University of Manitoba.

Additionally, M.W.K. was supported by the Natural Sciences and Engineering Research Council (NSERC) of Canada through the VADA Grant. We thank the participants, their parents, and caregivers, as well as the Misericordia Health Center. The graphical abstract was created using the BioRender license held by P.C. The authors also extend their gratitude to Drs. Nisha Singh, Ryan Cunnington, and Ankita Vaishampayan of The Chelikani Lab for their invaluable assistance and logistical support. Furthermore, we are grateful to the Manitoba Centre for Health Policy (MCHP) for providing socioeconomic data based on the postal codes.

