## Supplementary file for "Integrative analysis of taste genetics and the dental plaque microbiome in early childhood caries"

### Inclusion criteria for genes

The inclusion criteria for selecting the candidate genes were based on their roles in taste perception. This included 25 genes for bitter taste receptors, collectively known as the TAS2R family, which are part of the G-protein-coupled receptor (GPCR) superfamily and activated by bitter compounds. The other group of genes included the TAS1R family, which detect sweet and umami tastes through GPCR signaling. Additionally, several GPCRs involved in various taste signaling pathways were also included. The study also included genes encoding free fatty acid receptors, ion channels, and transporters for sodium balance, carbonic anhydrase genes involved in taste perception and pH regulation, and various other signal transduction proteins. A detailed list of these genes is provided in Supplementary **Table 1**. In our study, we utilized deep sequencing of gene exomes to identify novel variants, specifically frameshifts caused by insertions and deletions. However, no novel variants were found to be significantly associated with ECC in our analyses.

### Covariates included in the analyses

In addition to age and sex, the SEFI score and rural-urban information were included as confounders because of their association with ECC outcomes (Khan et al. 2024). To address potential biases related to multi-ancestry in our samples due to diverse ethnic groups and immigration patterns over time, we also incorporated the top five principal components (PCs) (**Figure S2A**). Although sequencing was conducted at multiple time points, batch information was not included as a covariate because most batch effects were captured by PC1, which showed a Spearman correlation of 0.81. Consequently, PC1 in our covariates effectively served as a proxy for batch effects (**Figure S2B**). Furthermore, including batch effects as a covariate did not improve the model. A comparison of genomic inflation factors indicated that adding batch effects after incorporating PCs worsened the model by increasing genomic inflation (**Figure S3**). This likely reflects the bias introduced by the strong correlation between PC1 and the batch effects. This observation was further supported by the Akaike information criterion (AIC) and Bayesian information criterion (BIC) values from covariate comparisons for individual models of the top variants (**Figure S4**).

**Table S1: List of the genes considered as candidate genes for targeted sequencing.**

| Gene Symbol | Function/Role | Type |
|---|---|---|
| ***TAS2R1 TAS2R3 TAS2R4 TAS2R5 TAS2R7 TAS2R8 TAS2R9 TAS2R10 TAS2R13 TAS2R14 TAS2R16 TAS2R19 TAS2R20 TAS2R30 TAS2R31 TAS2R38 TAS2R39 TAS2R40 TAS2R41 TAS2R42 TAS2R43 TAS2R45 TAS2R46 TAS2R50 TAS2R60*** | Bitter taste receptors | Taste G-protein-coupled receptors (GPCRs) |
| ***TAS1R1 TAS1R2 TAS1R3*** | Sweet and umami taste receptors | Taste GPCRs |
| ***SCNN1A SCNN1B SCNN1D SCNN1G*** | Sodium ion transport | Epithelial Sodium Channels (ENaC) |
| ***GPR161 GPR107 GPR87 ADGRG1 ADGRG6*** | Signal transduction and cellular communication | GPCRs |
| ***FFAR1 FFAR2 FFAR3 FFAR4*** | Free fatty acid receptors | GPCRs - Fatty Acid Receptors |
| ***GNAI2 GNAT3 GNAS GNAQ*** | Intracellular signaling via G proteins | G-proteins |
| ***PLCB2*** | Secondary messenger generation (phosphoinositide pathway) | Phospholipase C |
| ***CA6 CA7*** | pH regulation through bicarbonate production | Carbonic Anhydrases |
| ***RALB RAC1*** | Small GTPase-mediated signaling | Small GTPases |
| ***OTOP1*** | Proton transport and pH sensing in taste | Ion Channel |
| ***FZD6*** | Wnt signaling | Frizzled class receptor |
| ***GABBR1*** | Synaptic inhibition via GABA signaling | GABA Receptor |

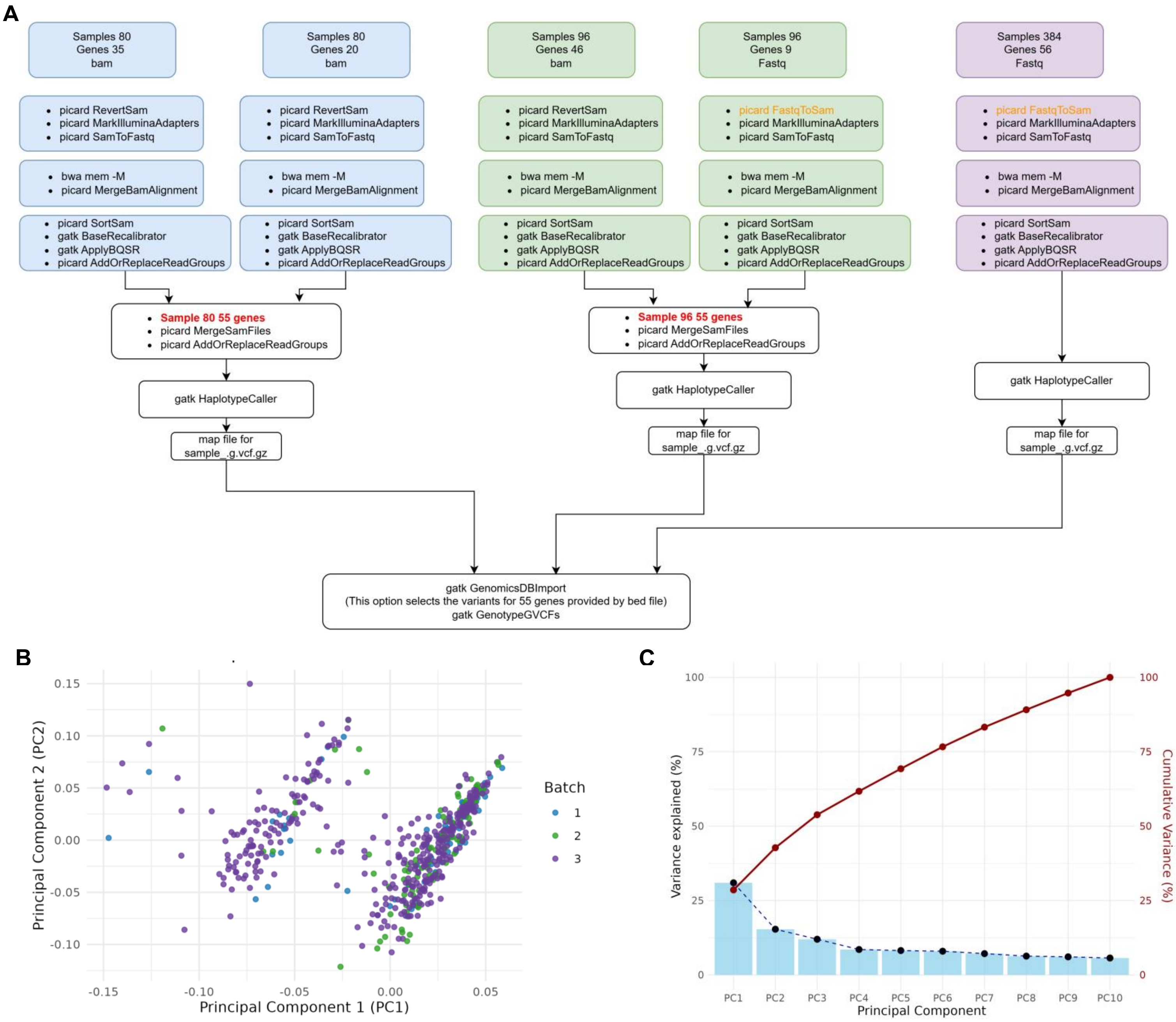


**Figure S1: Batch processing, batch effects, covariates, and principal components from genetic data.** (A) Flowchart of the steps used to combine data from different sequencing runs. Initially, 560 samples were run, but only 538 were retained after removing duplicated samples and age selection criteria of less than 72 months. (B) Batch effects in the quality-filtered genetic variants after selection cutoff, as explained in the Methods section. (C) Scree plot of principal components (PCs) showing the variance explained by individual components (left y-axis) and line plot of cumulative variance (right y-axis).

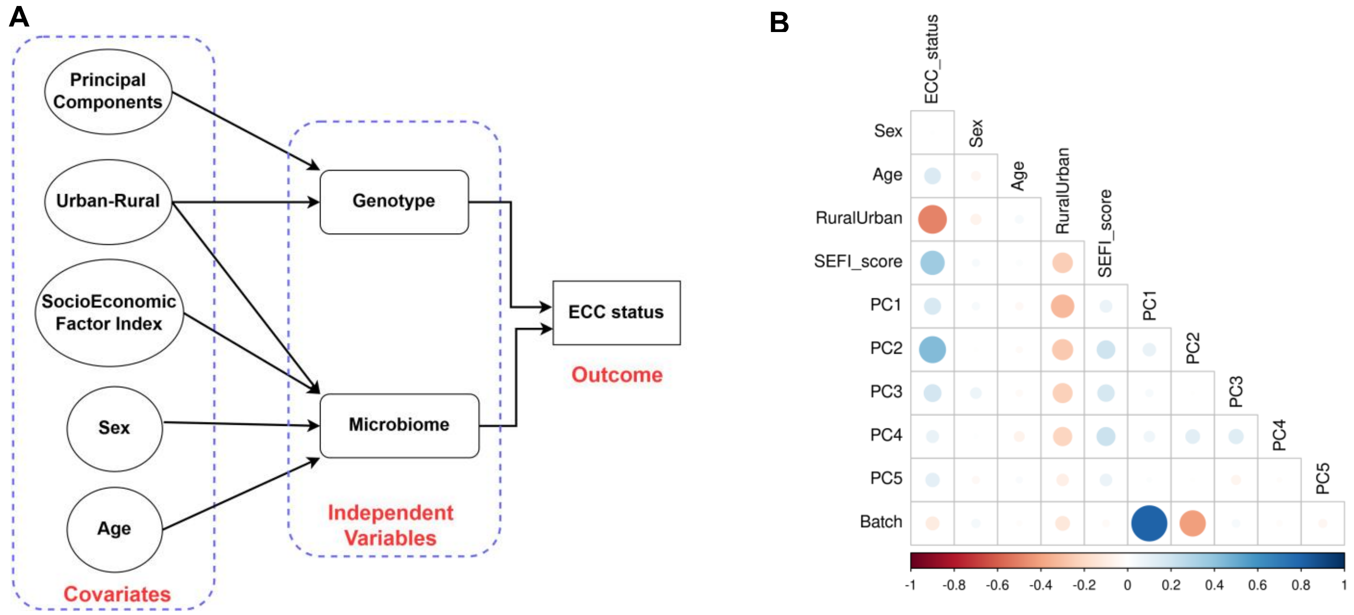


**Figure S2: Covariates used in this study.**
(A) Directed acyclic graph illustrating the hypothesized relationships between covariates, independent variables, and outcome in our study
(B) Spearman correlation between the covariates and the top five principal components (PCs) from the genetic data used in the study.

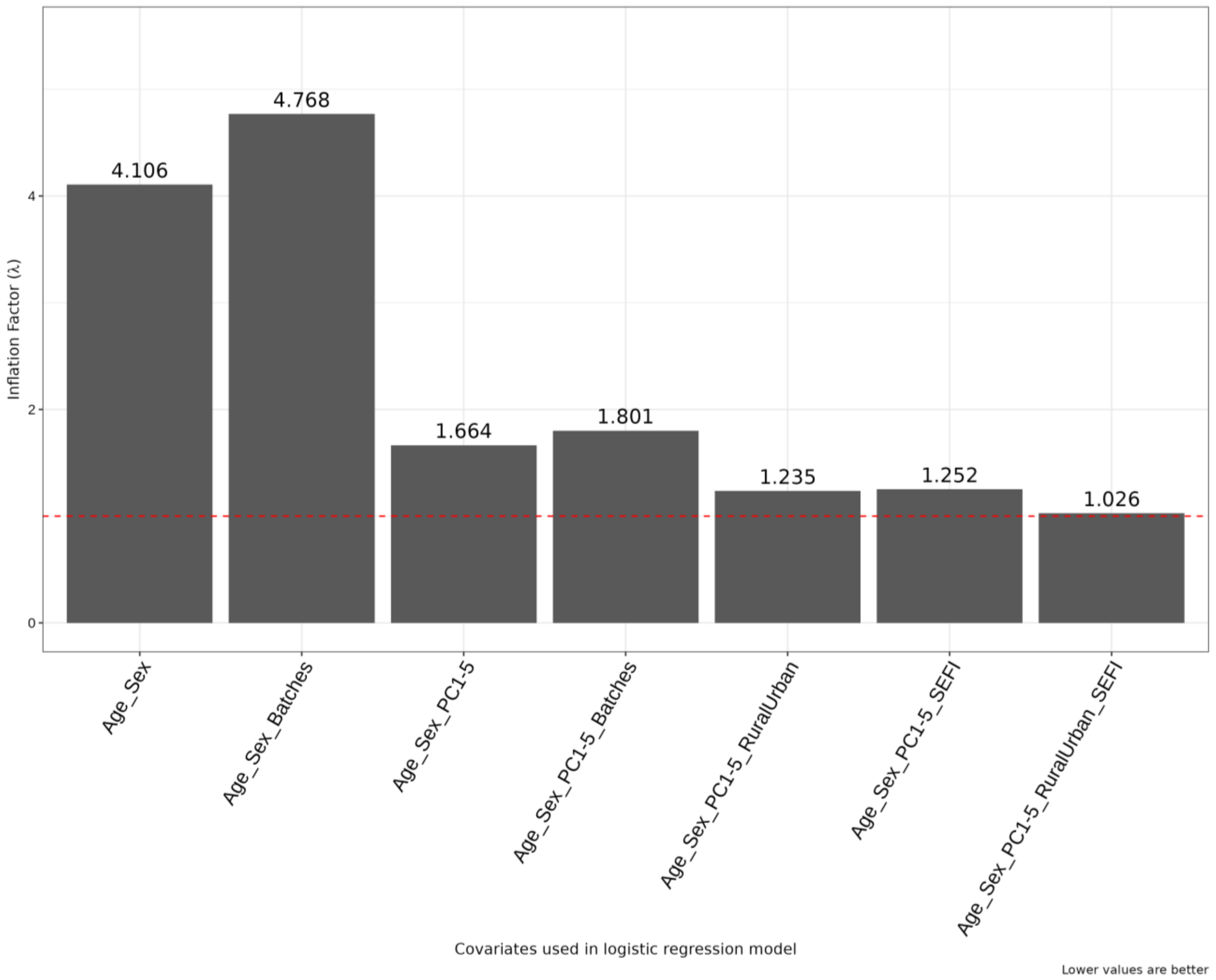


**Figure S3. Genomic inflation rate 'lambda'(λ) for *p*-values for all variants using different covariates.** The λ is a measure of the deviation of *p*-values from the null distribution, with λ = 1 indicating no inflation. Each bar represents the combinations of covariates, which include age, sex, batches, rural-urban status, SEFI score, and top 5 principal components (PC1-5).

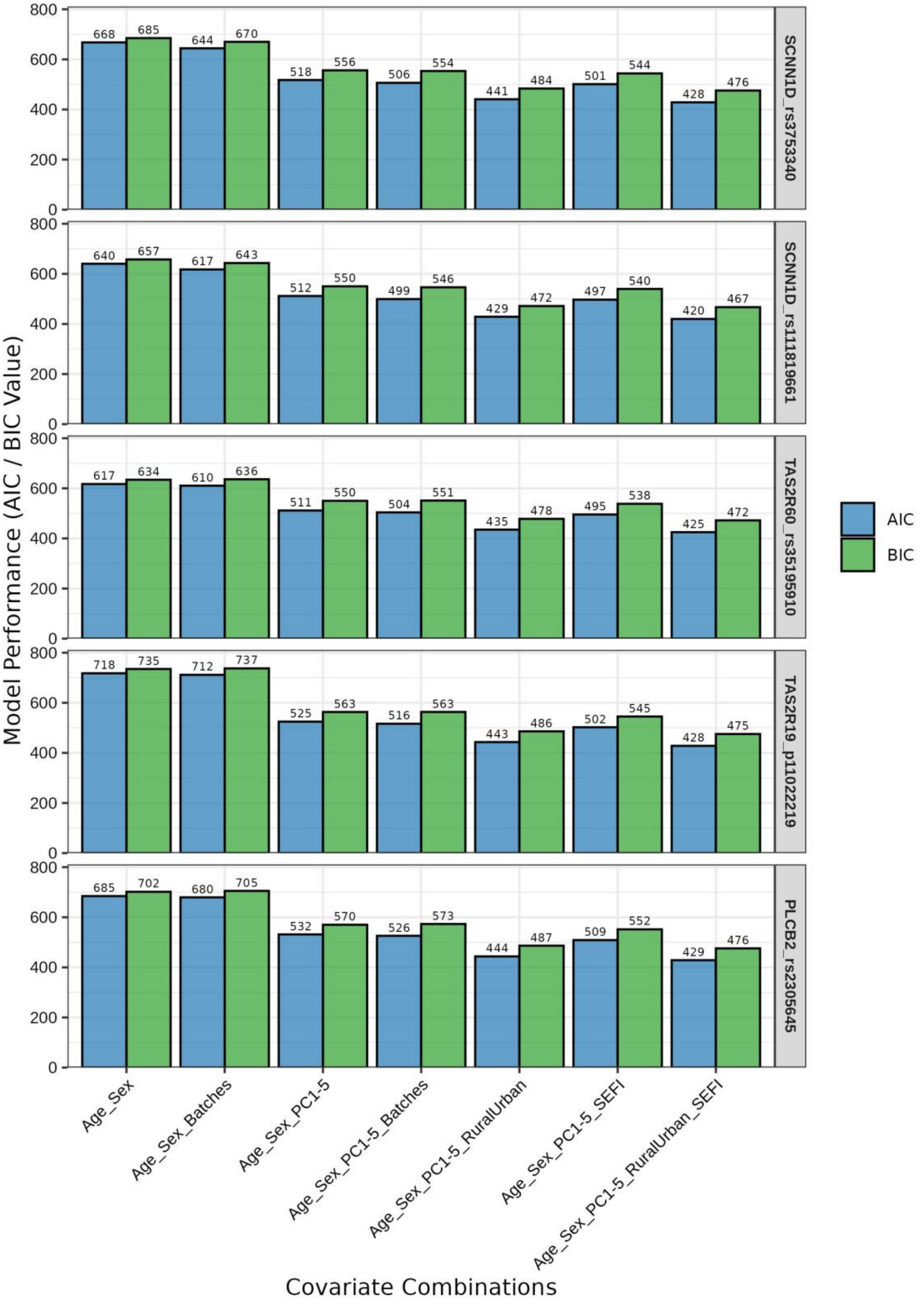


**Figure S4: Model evaluation for different covariates using the top five ECC-associated taste genetic variants.**

The top five variants identified using the base model for the association between variants and ECC status using age and sex as covariates. Various models were evaluated for combinations of covariates, including batches, rural-urban status, SEFI score, and the top five principal components (PC1-5). The *glm* model was used with ECC status as the outcome, with each variant as an independent variable, and the Akaike

Information Criterion (AIC) and Bayesian Information Criterion (BIC) values were compared. (The lower values are better).

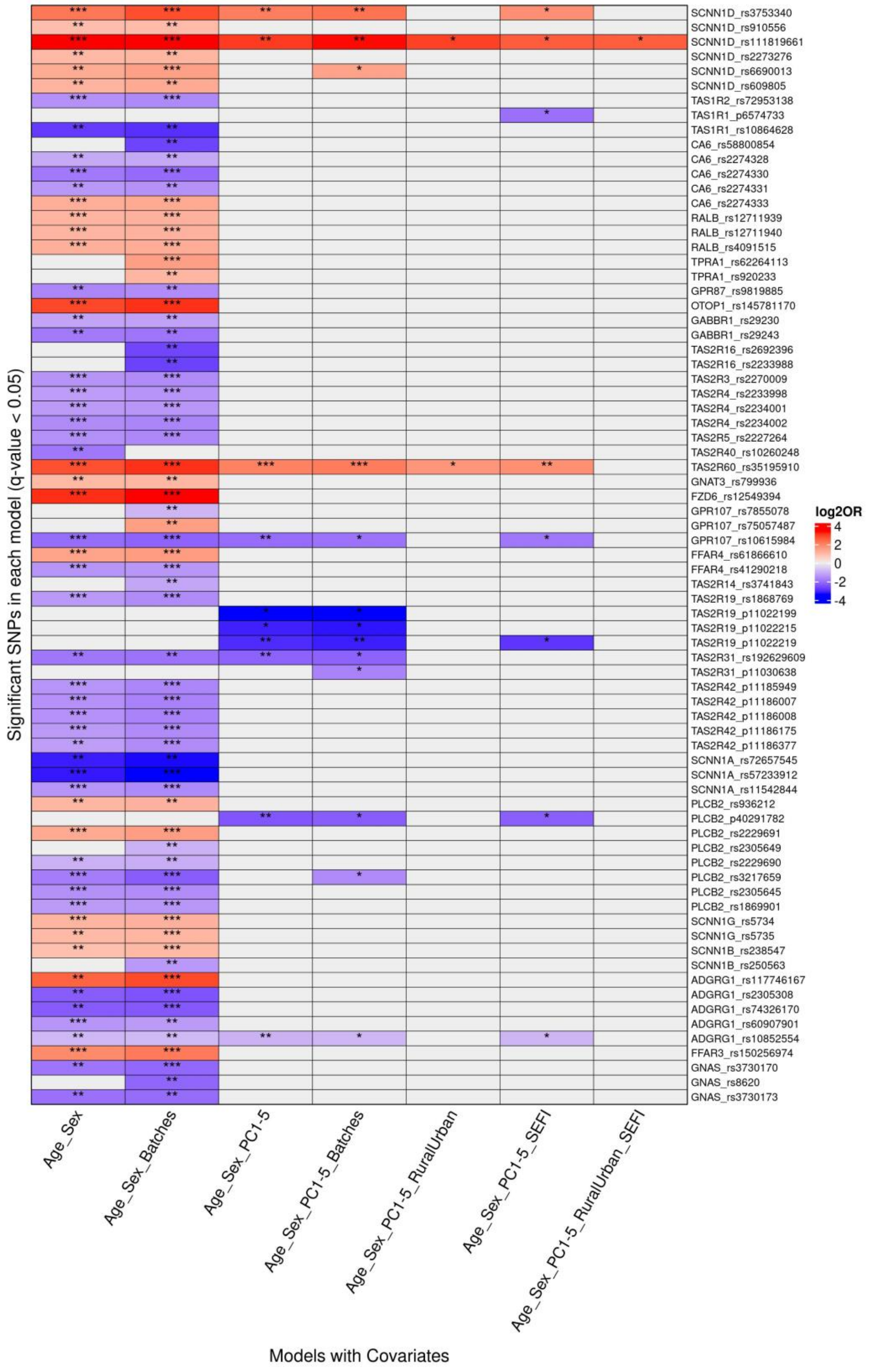


**Figure S5. Significant genetic variants across different association models for ECC outcome.**

Each row represents a specific variant identified by its chromosomal location and associated gene. The columns depict different statistical models based on covariate inclusion, which is given on the x-axis and includes a combination of age, sex, batch effects, rural-urban status, SEFI score, and the top five principal components from genetic variants in the 55 taste-associated candidate genes. The color represents the log2odds ratio, while asterisks denote levels of statistical significance (* $p < 0.05$, ** $p < 10^{-3}$, *** $p < 10^{-5}$) with white indicating no significant association.

polymorphism (SNP) that was tested in the analysis. The red dashed line represents the null expectation, where the observed values equal the expected values.
(B) Volcano plot. The x-axis represents the log2 odds ratio (OR), and the y-axis shows the -log10(*p*-value). Each point represents a variant, with the shape indicating the variant type. Red points denote significant associations for *q* <0.05, where *q*-values are BH adjusted *p*-value <0.05.
(C) Similar to A, but for the recessive genetic model.
(D) Similar to B, but for the recessive genetic model.
The Q-Q and volcano plots are from logistic regression models in PLINK for 55 taste-associated candidate genes, with age, sex, rural-urban status, SEFI score, and PCs 1–5 as covariates.
(E) Gene-wise association for rare, minimum allele frequency (MAF) < 0.01) and common variants (MAF > 0.01) with default weights in the SKAT method in R. The bar plot represents the *p*-values and *q*-values for the association between variants within each gene after removing singletons. The analysis was performed in the SKAT R-package using the SKAT-O option for optimal association with age, sex, rural-urban status, SEFI score, and top 5 principal components as covariates. The horizontal line indicates the significance value cutoff of <0.05.

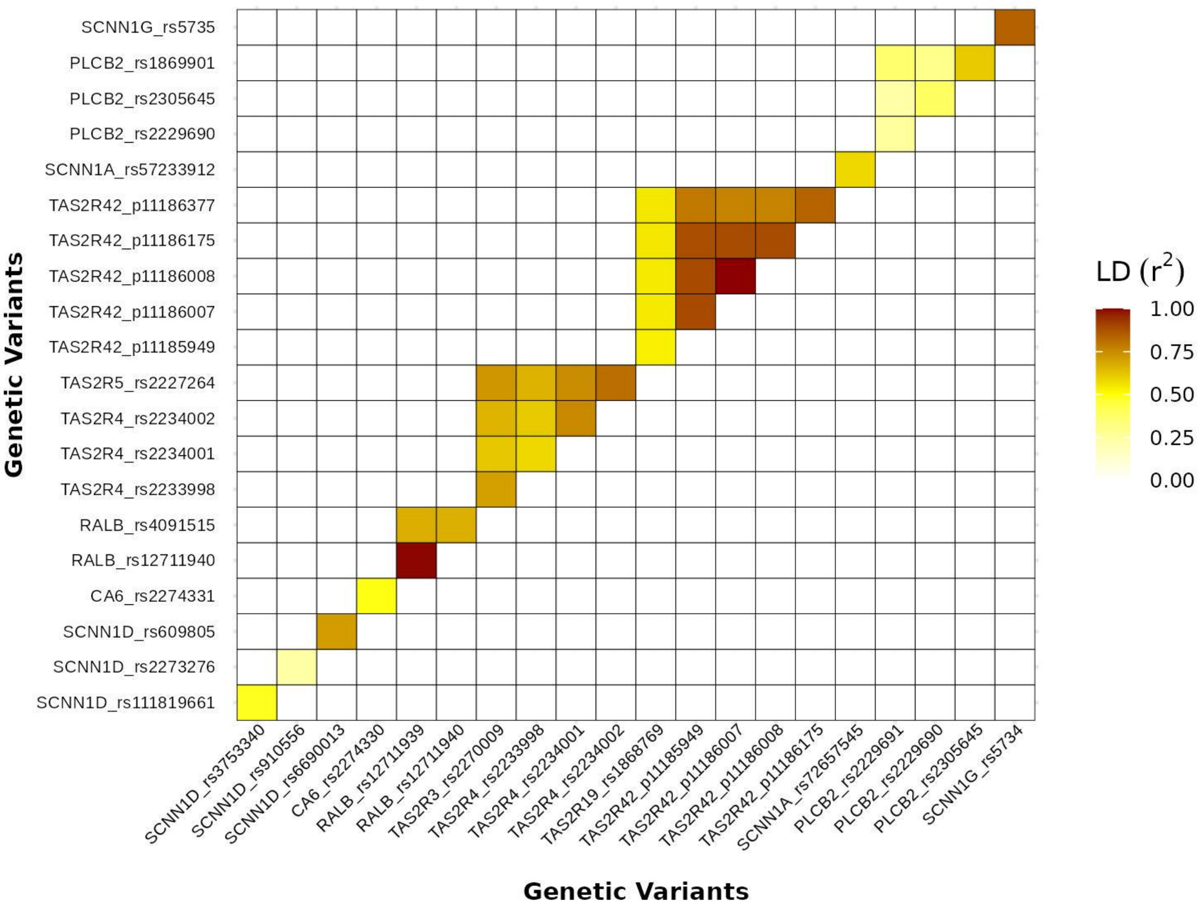


**Figure S7: Linkage disequilibrium (LD) analysis of the variants identified in the model with age and sex as covariates.**

Only the variants in *TAS2R3*, *TAS2R4*, and *TAS2R5* exhibited a moderate level of intergenic LD, whereas no significant intergenic LD was observed among the other variants.

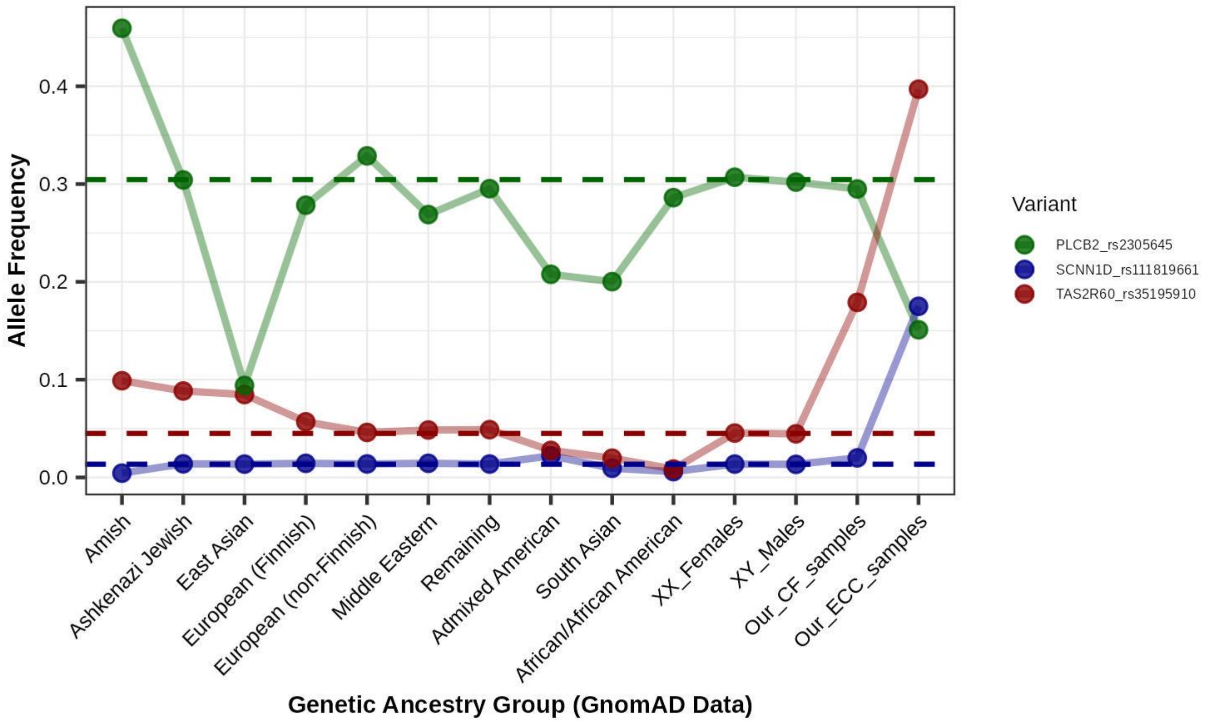


**Figure S8: Allele frequencies of ECC-associated significant taste-related genetic variants.**
The figure presents the allele frequencies of *SCNN1D*-rs111819661, *TAS2R60*-rs35195910, and *PLCB2*-rs2305645 across different genetic ancestry groups from the gnomAD database v4.1. Horizontal lines indicate the average allele frequency of the population for each variant.

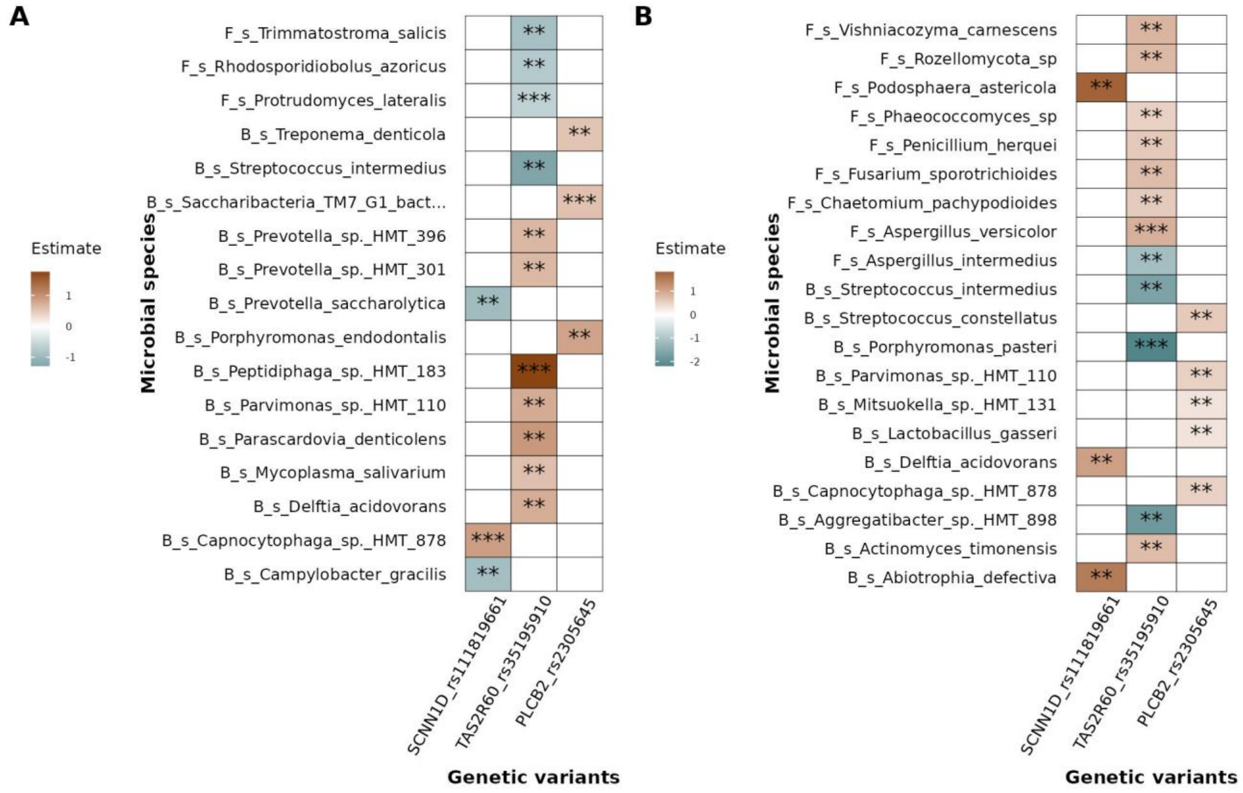


**Figure S9: Association between ECC-associated variants and microbial taxa in ECC and CF stratified samples.**

The heatmap illustrates the association between ECC-associated genetic variants and microbial taxa when adjusted for covariates such as age, sex, urban-rural status, SEFI score, and the top five principal components from the genetic data. The forest plot shows the coefficient of association from the MaAsLin2 method between the selected microbial species in the heatmap and ECC outcome. Positive coefficients favor ECC, whereas negative coefficients favor CF status. The analysis was performed for significant variants identified in the dominant genetic model and all microbial species with CLR normalization using a linear model. The heatmap value signifies the estimates values, with red showing a positive association between variants with alternate alleles and green representing a negative association. The *p*-values are (* $p < 0.05$, ** $p < 0.01$, *** $p < 0.001$). Microbial features are labeled as ‘B’ for bacterial and ‘F’ for fungal taxa. The prefix ‘s’ signifies species level, and ‘g’ indicates genus level. (A) Genus level (B) Species level.

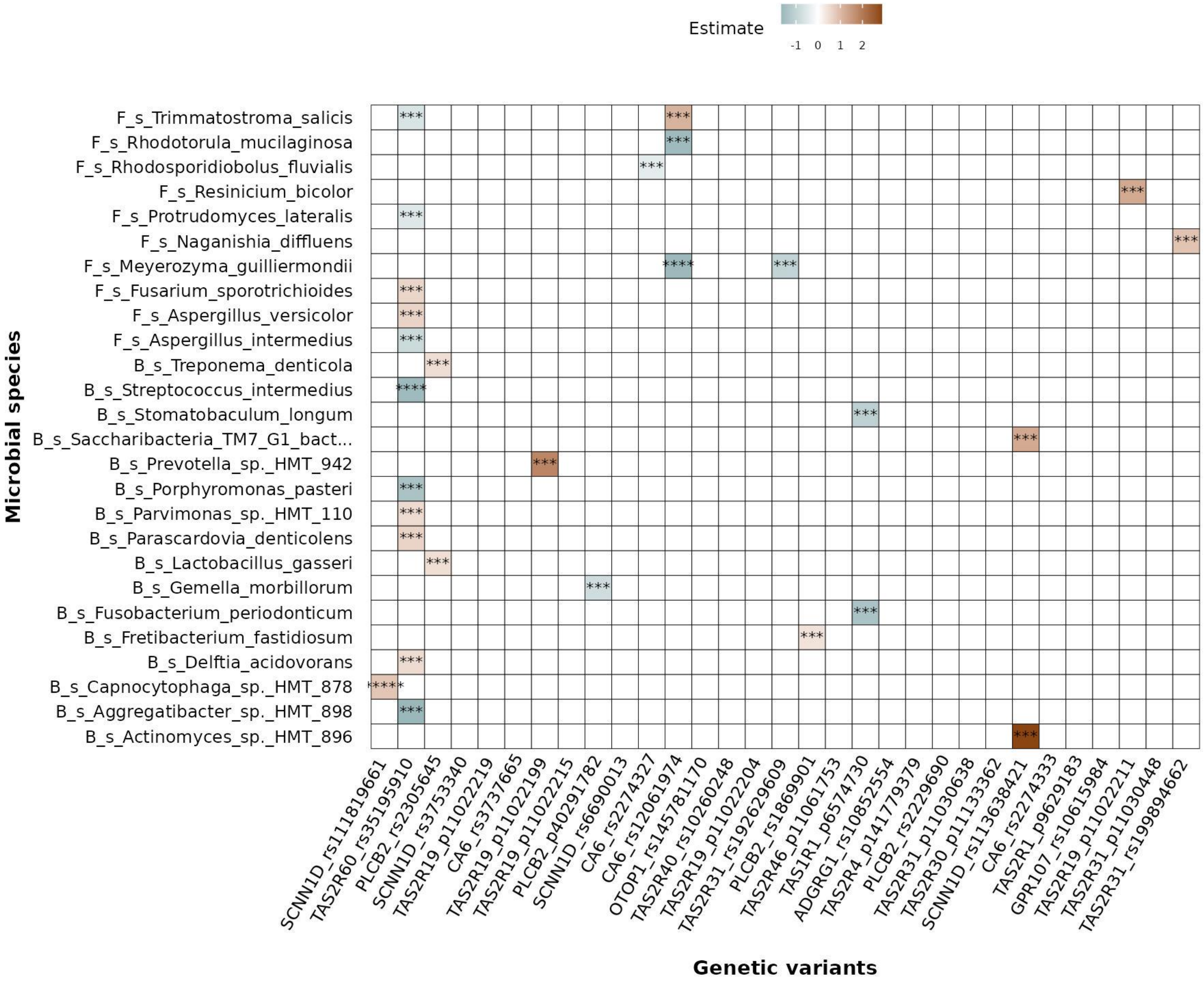


**Figure S10: Association between 31 top variants (5% of all quality-filter passing variants) based on *p* values for association with ECC and all microbial taxa.**

With covariates age, sex, rural-urban status, SEFI score, top 5 principal components, and ECC status The analysis was performed for significant variants identified in the dominant genetic model and all microbial species with CLR normalization using a linear model. The heatmap value signifies the estimates values, with red showing a positive association between the variant with the alternate allele and blue representing a negative association. The P-values are (* $p < 0.05$, ** $p < 0.01$, *** $p < 0.001$, **** $p < 0.0001$). Microbial features are labeled as ‘B’ for bacterial and ‘F’ for fungal taxa. The prefix ‘s’ signifies species level, and ‘g’ indicates genus level. Genetic variants were named using the Gene_rsID (Reference SNP ID) approach when rsID based on dbSNP138 was available. For variants lacking rsID, the naming convention Gene_p followed by the chromosomal position was applied, indicating novel variants not assigned in the dbSNP138 database.

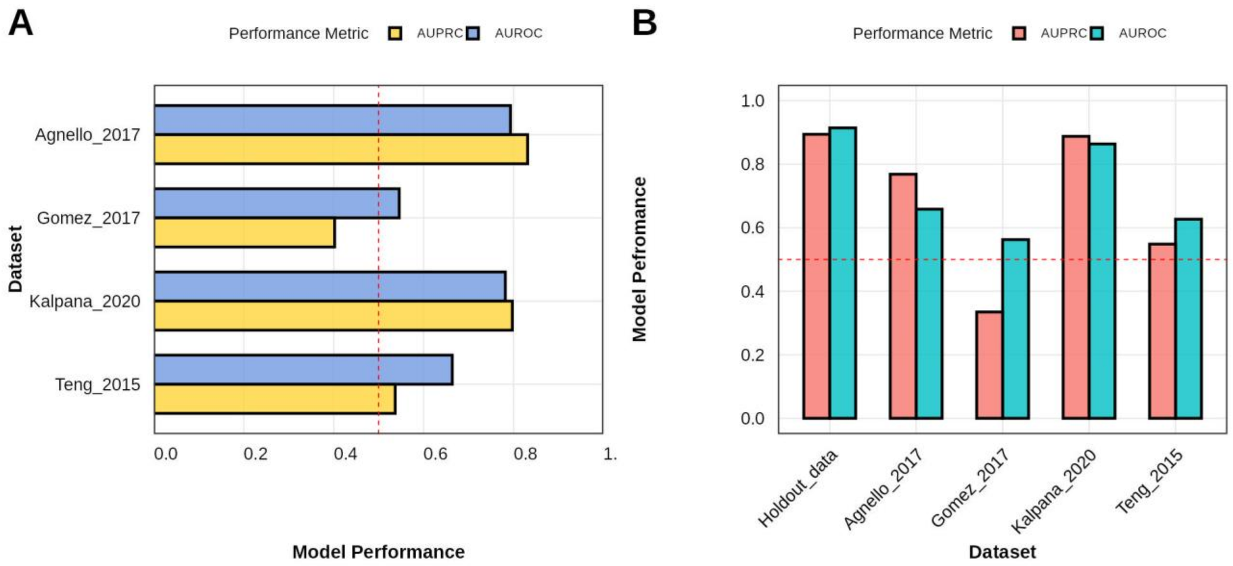


**Figure S11: Machine learning based classification for ECC status using microbiome data on external datasets**

(A) RF model performance on external datasets (without batch correction). Four external datasets were chosen to assess the RF model based on only microbiome data, and the model performance was assessed using AUROC and AUPRC values.

(B) ECC prediction model performance on external datasets (without batch correction) based on the MRS score on CLR-normalized datasets.
